# Missense variants in human ACE2 modify binding to SARS-CoV-2 Spike

**DOI:** 10.1101/2021.05.21.445118

**Authors:** Stuart A. MacGowan, Michael I. Barton, Mikhail Kutuzov, Omer Dushek, P. Anton van der Merwe, Geoffrey J. Barton

**Affiliations:** Division of Computational Biology, School of Life Sciences, University of Dundee, Dow Street, Dundee, DD1 5EH, Scotland, UK; Sir William Dunn School of Pathology, University of Oxford, Oxford, Oxfordshire, United Kingdom

## Abstract

SARS-CoV-2 infection begins with the interaction of the SARS-CoV-2 Spike (Spike) and human angiotensin-converting enzyme 2 (ACE2). To explore whether population variants in ACE2 might influence Spike binding and hence infection, we selected 10 ACE2 variants based on affinity predictions and prevalence in gnomAD and measured their affinities for Spike receptor binding domain through surface plasmon resonance (SPR). We discovered variants that enhance and reduce binding, including two variants with distinct population distributions that enhanced affinity for Spike. ACE2 p.Ser19Pro (ΔΔG = ± 0.59 0.08 kcal mol^−1^) is often seen in the gnomAD African cohort (AF = 0.003) whilst p.Lys26Arg (ΔΔG = 0.26 ± 0.09 kcal mol^−1^) is predominant in the Ashkenazi Jewish (AF = 0.01) and European non-Finnish (AF = 0.006) cohorts. Carriers of these alleles may be more susceptible to infection or severe disease and these variants may influence the global epidemiology of Covid-19. We also identified three rare ACE2 variants that strongly inhibited (p.Glu37Lys, ΔΔG = −1.33 ± 0.15 kcal mol^−1^ and p.Gly352Val, predicted ΔΔG = −1.17 kcal mol^−1^) or abolished (p.Asp355Asn) Spike binding. These variants may confer resistance to infection. Finally, we calibrated the mCSM-PPI2 ΔΔG prediction algorithm against our SPR data, give new predictions for all possible ACE2 missense variants at the Spike interface and estimate the overall burden of ACE2 variants on Covid-19 phenotypes.

## 1 Introduction

The COVID-19 pandemic is one of the greatest global health challenges of modern times. Although the disease caused by the severe acute respiratory syndrome coronavirus 2 (SARS-CoV-2) is usually cleared following mild symptoms, it can progress to serious illness and death^1^. Besides the clear risks associated with age and comorbidities^2,3^, there could be a genetic component that predisposes some individuals to worse outcomes^4^. Genetic association studies have already identified several loci involved in Covid-19 risk^5^. Identifying further genetic factors of COVID-19 susceptibility has implications for clinical decision making and epidemic dynamics. Genetic variation may constitute hidden risk factors and, in some cases, explain why otherwise healthy individuals in low-risk groups experience severe disease. The identification of specific genetic variants that influence the severity and progression of COVID-19 presents the opportunity for predictive diagnostics, early intervention and personalised treatments whilst the population distribution of such variants could contribute to population specific risk.

Human angiotensin-converting enzyme 2 (ACE2) is the host cell receptor that SARS-CoV-2 exploits to infect human cells^1,6^. As this is the same receptor used by the SARS coronavirus (SARS-CoV) that caused the SARS outbreak in 2002, the detailed body of knowledge built around SARS-CoV infection is relevant to understanding SARS-CoV-2^1,6,7^. The spike glycoprotein (Spike) is the coronavirus entity that recognises and binds host ACE2. Both SARS coronavirus Spikes include an S1 domain that contains ACE2 recognition elements and an S2 domain that is responsible for membrane fusion^6^. Spike is primed for cell fusion by cleavage with host furin^8^ and TMPRSS2^6^, in SARS-CoV cleavage by TMPRSS2 is thought to be promoted upon formation of the ACE2 Spike complex^9^. The S1 receptor binding domains (RBDs) from both SARS-CoV^10^ and SARS-CoV-2^11^ have been co-crystallised with human ACE2. The RBDs from both viruses are similar in overall architecture and interface with roughly the same surface on ACE2. Differences are apparent in the so-called receptor binding motif, which is the region of the RBD responsible for host range and specificity of coronaviruses^10–12^. The binding affinity of Spike and ACE2 is known to be correlated to the infectivity of SARS-CoV and is determined by the complementarity of the interfaces^10,12^. However, despite its essential role in infection, risk variants in ACE2 have not been conclusively identified in genetic association studies.

Missense variants located in protein-protein interaction interfaces can affect altered binding characteristics^13,14^ and in the context of virus-host interactions, have been shown to effect susceptibility^15^. This indicates the potential of missense variants in ACE2 to alter Spike binding and therefore influence a key step in SARS-CoV-2 infection. This is also suggested by the fact that the host range of coronaviruses is partly determined by the complementarity of the Spike receptor binding motif and the target hosts’ ACE2 sequence^10,12^. A few studies^4,16–21^ have addressed this question and gave rise to conflicting conclusions regarding the effects of specific variants on the interaction affinity and their relevance to the pandemic generally. The strongest of these used data from a published deep mutagenesis binding screen^22^ to assess the effect of ACE2 population variants on Spike affinity and confirmed the effects of five key variants with further biochemical assays^17^. In our own previous contribution^23^, we employed the mCSM-PPI2 protein-protein interaction affinity prediction algorithm^13^ to assess the effects of ACE2 variants on the binding of SARS-CoV-2 Spike and predicted that three reported ACE2 variants would strongly inhibit or abolish binding (p.Asp355Asn, p.Glu37Lys and p.Gly352Val). Confidence was given to these predictions by the performance of mCSM-PPI2^13^ in comparison to binding data for 26 ACE2 mutants in complex with SARS-CoV Spike RBD^12^. Here we report experimental binding affinities of 10 ACE2 variants for isolated SARS-CoV-2 Spike RBD determined via surface plasmon resonance (SPR). These results give better insight into the effect of ACE2 variants on SARS-CoV-2 Spike binding, revealing additional variants that enhance Spike binding and also test the quality of our predictions. The SPR data also allowed us to recalibrate the mCSM-PPI2 predictions to provide more accurate estimates of the effect of interface variants that we did not test experimentally.

## 2 Results

### 2.1 ACE2 variant affinities for SARS-CoV-2 Spike

Figure 1 highlights the mutated residues on the structure of ACE2^11^ in complex with SARS-CoV-2 Spike. We determined the binding affinity of 10 ACE2 mutants for isolated SARS-CoV-2 Spike RBD via surface plasmon resonance (SPR) to identify variants that may influence an individual’s response to infection. Nine of these mutants were selected from the 241 ACE2 missense variants reported in gnomAD^24^ on the basis of our previous computational predictions^23^ and their reported prevalence, whilst the tenth mutant (p.Thr27Arg) was predicted to enhance Spike binding more than any other possible mutation at the interface.

**Figure 1.**
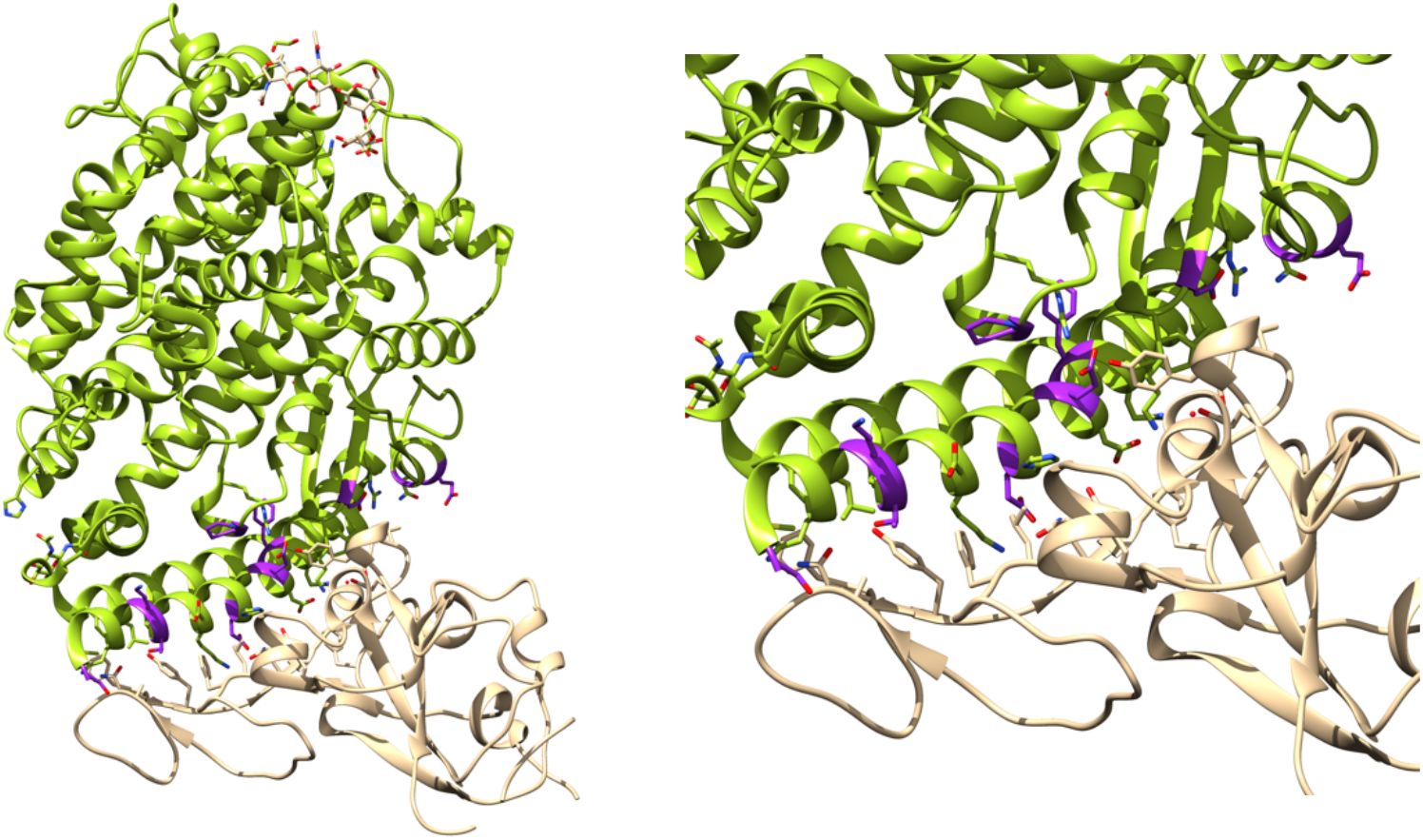
*Binding affinity determination of ACE2 variants with SARS-CoV-2 Spike. Left: ACE2 (green) in complex with Spike RBD (tan) complex from biological assembly 1 derived from PDB ID: 6vw1^11^. The positions that were mutated in this work are highlighted magenta. Right. The ACE2 Spike interface. Figure generated with Jalview^25^ and UCSF Chimera^26^.*

Table 1 presents experimentally determined ΔΔG and mCSM-PPI2^13^ predictions for the 10 ACE2 mutants together with predicted data for a further three variants, alongside variant population frequencies and RBD interacting residues. Our SPR data were collected in two batches. The first batch comprised four variants, two were predicted to strongly reduce or abolish Spike binding (p.Glu37Lys and p.Asp355Asn) and two predicted to enhance Spike binding (p.Gly326Glu and Thr27Arg)^23^. The SPR measurements showed strongly reduced binding for p.Glu37Lys and the total abolition of binding for p.Asp355Asn (within the concentration range assayed), in agreement with the predictions. In contrast, p.Gly326Glu and Thr27Arg, which had been predicted to enhance binding, displayed decreased and slightly decreased binding, in disagreement with the predictions. These discrepancies motivated a second set of SPR measurements that included six of the most prevalent ACE2 variants close to the Spike binding site. Surprisingly, the two most common ACE2 variants tested bound SARS-CoV-2 Spike more strongly than reference ACE2. These were p.Lys26Arg (ΔΔG = 0.26 ± 0.09), which has the highest allele frequency of ACE2 variants near the Spike interface, and p.Ser19Pro (ΔΔG = 0.59 ± 0.09) that has the second highest frequency. Two other variants in this series also increased Spike binding (p.Phe40Leu and p.Pro389His) despite being over 8 Å from the closest Spike residue. Finally, p.Glu329Gly may cause a slight reduction in binding (ΔΔG = −0.09 ± 0.09) whilst p.Glu35Lys had an inhibitory effect (ΔΔG = −0.36 ± 0.09). These results show that ACE2 variants can both enhance and inhibit Spike binding, properties that may reasonably be associated with susceptibility and resistance phenotypes to SARS-CoV-2 infection.

**Table 1.**
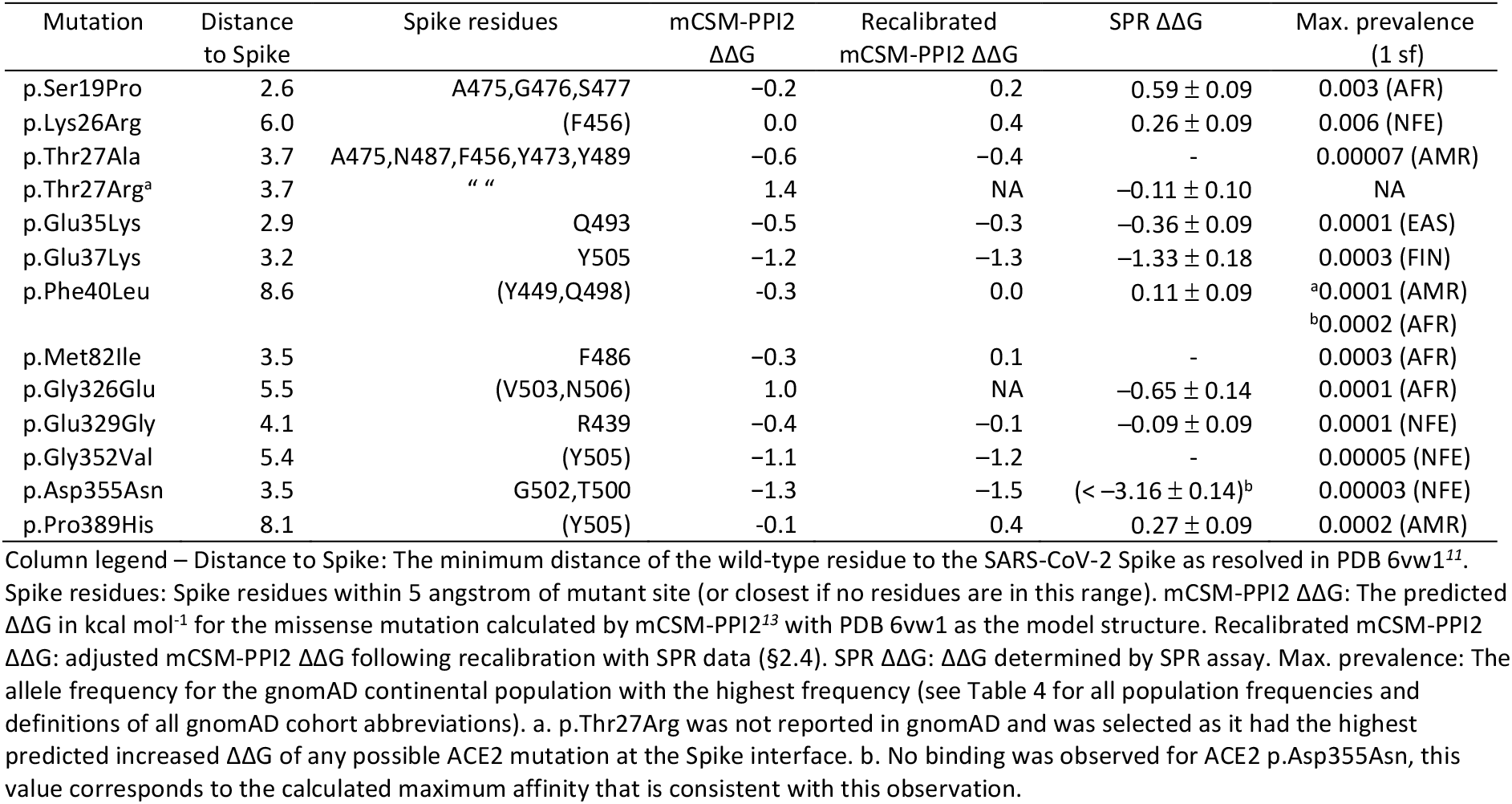
*Surface plasmon resonance derived ΔΔG and mCSM-PPI2^13^ predictions for ACE2 mutants, including gnomAD^24^ missense variants, at or near the ACE2 Spike interface. See Supplementary Table 1 for mCSM-PPI2 predictions for all gnomAD variants in the ACE2 ectodomain.*

### 2.2 Structural features of affinity modifying variants

Figure 2 and Figure 3 illustrate the mCSM-PPI2 provided models of p.Ser19Pro and p.Lys26Arg. p.Lys26Arg was not predicted to make any new well-defined contacts with Spike residues but it is predicted to adopt a conformation that extends toward Spike residues Y473 and F456, coming within 7.8 Å and 6.1 Å of these residues’ aromatic rings, respectively, slightly beyond typical amino-aromatic interaction distances^27^ but potentially favourable when dynamics are considered. Similarly, p.Ser19Pro also does not introduce any new Spike contacts according to the mCSM-PPI2 model and so the enhanced affinity is difficult to explain but it is has been suggested that the Pro mutant stabilises the helix to favour Spike interaction^22^. ACE2 p.Ser19Pro is of further interest because of its proximity to Spike Ser477, which has mutated to Asn in circulating SARS-CoV-2 strains and these ACE2 and RBD variants have been found to interact^28^. The mCSM-PPI2 structural models of the p.Asp355Asn, p.Glu37Lys and p.Gly352Val were discussed in detail in our previous work^23^. In summary, p.Asp355Asn was predicted to introduce a number of steric clashes at the interface, whilst p.Glu37Lys abolished an H-bond between ACE2 Glu37 and RBD Tyr505.

**Figure 2.**
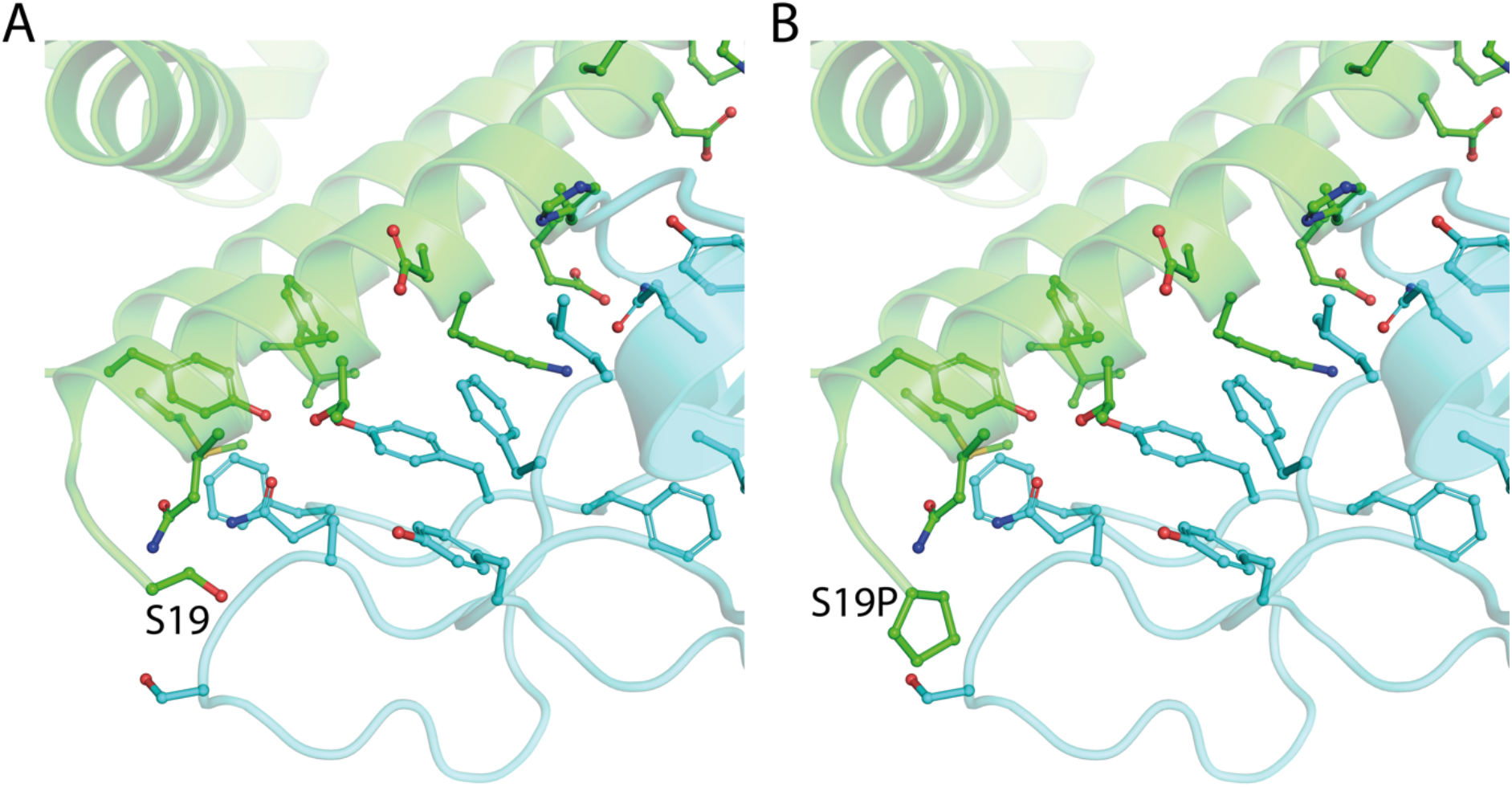
*The structure of ACE2 (green) gnomAD^24^ missense variant p.Ser19Pro that enhances Spike (light blue) binding affinity. A. The environment of ACE2 Ser19 from PDB ID: 6vw1^11^. B. Model of ACE2 p.Ser19Pro in complex with Spike. The mutant structure was modelled onto 6vw1 with mCSM-PPI13. Figure created with PyMol^29^.*

**Figure 3.**
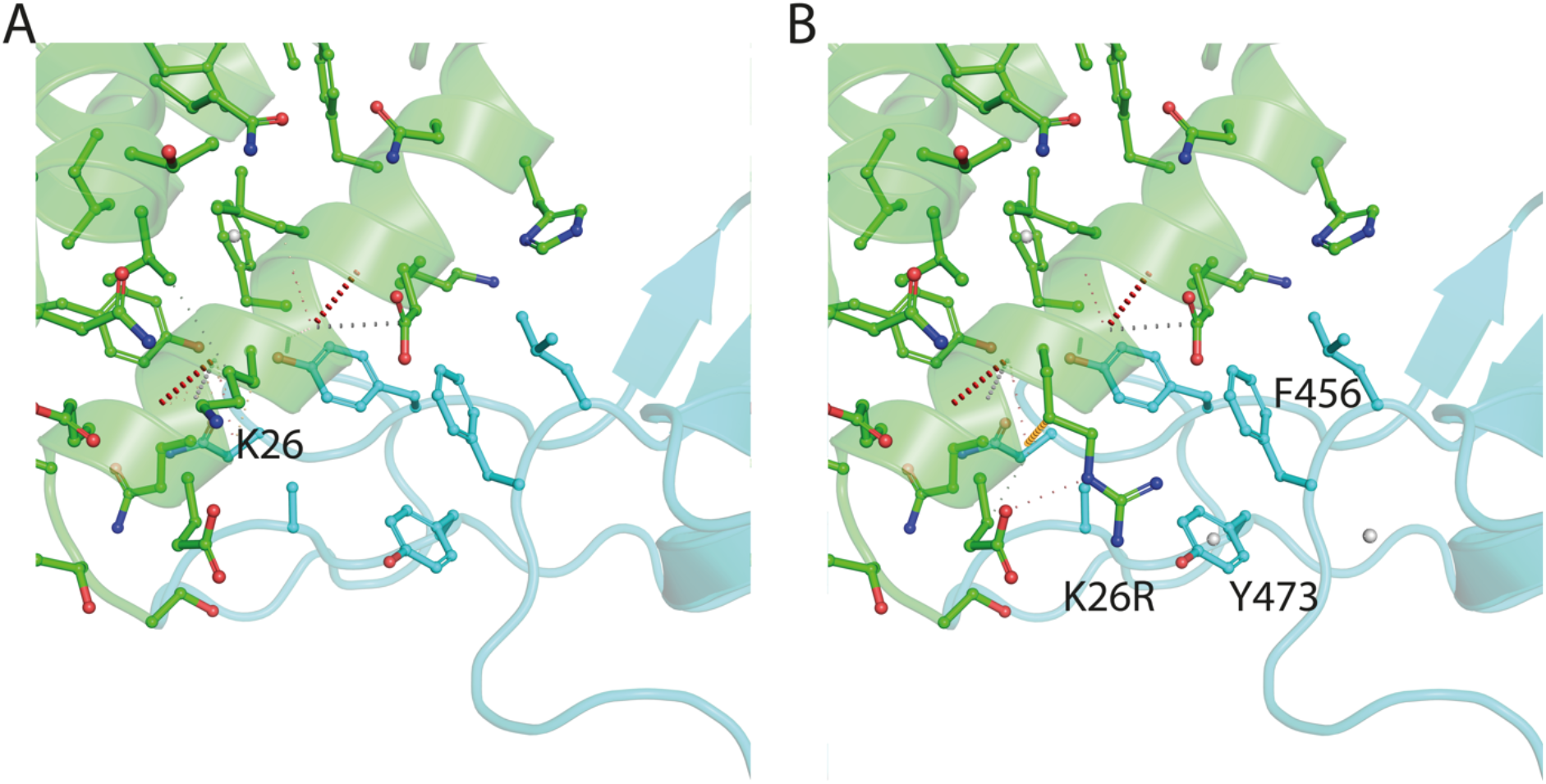
*The structure of ACE2 (green) gnomAD^24^ missense variant p.Lys26Arg that enhances Spike (light blue) binding affinity. A. The environment of ACE2 Lys26 from PDB ID: 6vw1^11^. B. Model of ACE2 p.Lys26Arg in complex with Spike. The mutant structure was modelled onto 6vw1 with mCSM-PPI13. Figure created with PyMol^29^.*

### 2.3 Enhanced Spike binding ACE2 variants are relatively common in European (p.Lys26Arg) and African/African-American (p.Ser19Pro) populations

The enhanced binding by p.Lys26Arg and p.Ser19Pro is particularly interesting since this may effect carrier susceptibility or vulnerability toward SARS-CoV-2 infection and they have relatively high prevalence in the gnomAD^24^ populations. p.Lys26Arg is the most common missense variant in the ACE2 ectodomain (Total allele count = 797, Total allele frequency = 0.004) and is predominant in the Ashkenazi Jewish cohort (ASJ AF = 0.01) and the European (non-Finnish) population (NFE, AF = 0.006). Amongst NFE sub-populations, it is most prevalent in north-western Europeans (AF = 0.007) and least prevalent in southern Europeans (AF = 0.003) and Estonians (AF = 0.003). p.Lys26Arg was also observed within this frequency range in Latino/Admixed American (AMR, AF = 0.003) samples. The variant is less frequent in Finnish (FIN, AF = 0.0005), African/African-American (AFR, AF = 0.001), South Asian (0.001) and, especially, East Asian (0.00001) samples. Interestingly, gnomAD reports a second variant at this site, p.Lys26Glu, suggesting that the position is especially tolerant to mutation. p.Ser19Pro is the next most common ACE2 missense variant in proximity to the Spike binding site (AC = 64, AF = 0.0003) and has the highest positive ΔΔG of all those tested (ΔΔG = 0.59 ± 0.03). This variant is practically unique to the African/African-American gnomAD population (AFR, AC = 63, AF = 0.003). The only other observation of this variant in gnomAD is in a single heterozygote in the labelled “Other” cohort. As a result of these distinct population distributions, it is possible that these variants could contribute to some of the observed epidemiological^30^ variation between populations and ethnic groups.

Given the prevalence of these variants we checked for their occurrence in recent GWAS studies on Covid-19 related phenotypes. Table 2 presents GWAS association results for ACE2 p.Lys26Arg from the Covid-19 Host Genetics Initiative (HGI)^31^. These data show some consistency with the hypothesis that p.Lys26Arg contributes additional risk for more severe Covid outcomes, whilst not affecting the likelihood of infection. In the very severe respiratory confirmed Covid vs. population contrast (study A2), a non-statistically significant increased risk was reported (β = 0.38 ± 0.24, p = 0.12, p-het = 0.74). Other Covid HGI data summaries for contrasts testing for alleles associated with the risk of SARS-CoV-2 infection suggest the variant does not play a role in infection acquisition. Covid vs. lab/self-reported negative (β = −0.18 ± 0.14, p = 0.21, p-het = 0.92); Covid vs. population (β = −0.03 ± 0.11, p = 0.81, p-het = 0.65) and predicted Covid from self-reported symptoms vs. predicted or self-reported non-Covid (β = −0.10 ± 0.24, p = 0.67, p-het = 0.08). Although none of these tests achieved genome wide significance, it would be worthwhile to reassess the data after controlling for other loci with the greatest effect sizes, sex or other appropriate stratifications.

**Table 2:** *Covid Host Genetics Initiative summaries for p.Lys26Arg (rs4646116).*

| Covid HGI study and description | Beta | SE | p | p-het | meta N | meta AF | N Cases | N Controls |
| --- | --- | --- | --- | --- | --- | --- | --- | --- |
| A2 - very severe respiratory confirmed covid vs. population | **0.38** | 0.24 | 0.12 | 0.74 | 35,222 | 0.003 | 2,972 | 284,472 |
| B1 - hospitalized covid vs. not hospitalized covid |  |  | Variant not reported |  |  |  | 1,389 | 5,879 |
| B2 - hospitalized covid vs. population |  |  | Variant not reported |  |  |  | 6,492 | 1,012,809 |
| C1 - covid vs. lab/self-reported negative | **-0.18** | 0.14 | 0.21 | 0.92 | 64,950 | 0.004 | 11,181 | 116,456 |
| C2 - covid vs. population | -0.03 | 0.11 | 0.81 | 0.65 | 972,525 | 0.005 | 17,607 | 1,345,334 |
| D1 - predicted covid from self-reported symptoms vs. predicted or self-reported non-covid | -0.10 | 0.24 | 0.67 | 0.08 | 10,655 | 0.006 | 1,777 | 18,895 |

### 2.4 Recalibrated mCSM-PPI2 ΔΔG predictions

Our SPR data provide accurate readouts of the effect of ACE2 variants on Spike binding, but we could not feasibly carry out experiments for all possible ACE2 mutations at the interface. For variants we have not studied, predictions and other high-throughput datasets can be useful, so long as their applicability and limitations are well-understood. In our previous work^23^, we calibrated predictions against relative binding data for ACE2/SARS-CoV Spike, we now improve this by recalibrating the mCSM-PPI2 predictions with our ACE2/SARS-CoV-2 Spike SPR dataset.

Figure 4 compares experimental and mCSM-PPI2^13^ predicted ΔΔG. The eight mutants with predicted ΔΔG < 0 kcal mol^−1^ are linearly correlated with the experimental ΔΔG (R^2^ =0.91, p = 0.0006), whilst the two mutants with predicted ΔΔG > 1 kcal mol^−1^ (p.Gly326Glu and Thr27Arg) do not follow this behaviour. The absence of detectable binding for p.Asp355Asn and the strongly reduced binding observed for p.Glu37Lys, support our original determination that predicted ΔΔG < –1 kcal mol^−1^ was a reliable indicator of inhibitory variants. The difficulty mCSM-PPI2 has with affinity enhancing variants is not altogether surprising since our original calibration showed poorer performance for these variants^23^, which may be because there were fewer experimentally determined affinity enhancing variants in the algorithm’s training data. Structurally, although the mCSM-PPI2 model of p.Gly326Glu predicted new contacts with Spike^23^, it is possible that reduced torsional flexibility at mutant Glu326 has other effects that were not recognised (a similar argument applies for p.Ser19Pro but here the error is small). The subtler affinity changes measured for the five variants with predicted ΔΔG between −0.5 and 0.0 kcal mol^−1^, validate our original ambiguity towards those predictions, but the strong linear correlation within this narrow range is encouraging. Taken together, these observations suggest that predictions within the range of p.Glu36Lys (ΔΔG^pred^ =−1.2 kcal mol^−1^) and p.Lys26Arg (ΔΔG^pred^ =0.0 kcal mol^−1^) may be rescaled to provide a better estimate of actual ΔΔG. Recalibrated ΔΔG yields correct predictions for all eight mutations with negative predicted ΔΔG and measurable ΔΔG from SPR (Table 1). This lends additional confidence in our prediction that ACE2 p.Gly352Val strongly reduces Spike binding (in agreement with published deep mutagenesis experiments^22^).

**Figure 4.**
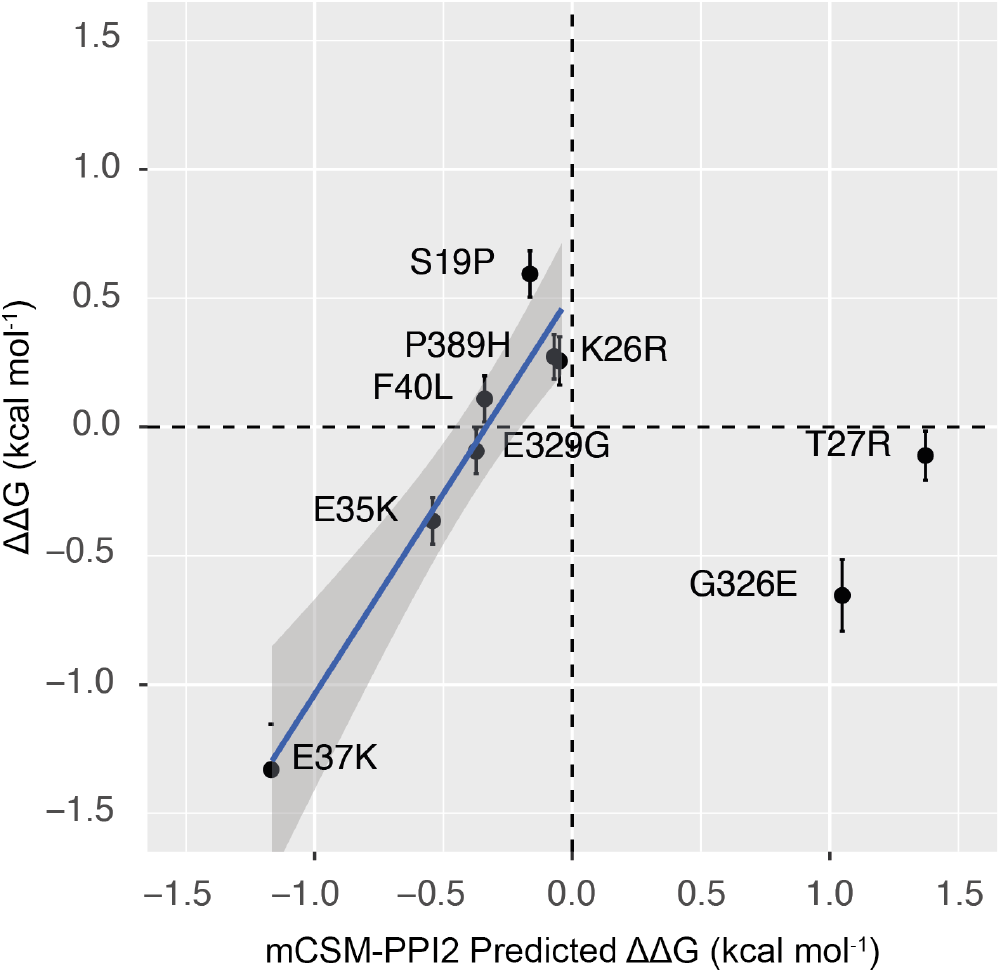
*Comparison of SPR ΔΔG with the mCSM-PPI2 prediction for 9 ACE2 mutations. The linear regression was fit to the eight ACE2 mutants with predicted ΔΔG < 0 kcal mol^−1^ (slope = 1.56, intercept = 0.52, R^2^ = 0.91, p = 0.0006). The shaded area indicates the 95 % confidence interval for predicted SPR ΔΔG from the linear model. Figure generated with R ggplot2.*

### 2.5 Predicted burden of rare ACE2 variants with Spike affinity phenotypes

Existing human variation datasets are well-powered to detect common variation (1KG was estimated to detect >99% SNPs with MAF >1%^32^ and gnomAD is substantially larger) in the sampled populations but they are far from comprehensive with respect to rare variation^24^. Rare variants in ACE2 that influence Spike binding could have implications for the epidemiology of COVID-19 in addition to the consequences for affected individuals. If there were 10 such variants with an allele frequency of 1 in 50,000, their collective occurrence might be as high as 1 in 5,000 (discounting linkage) and when this is considered alongside the possibility that a high proportion of the global population will be exposed to SARS-CoV-2 it becomes clear that such effects should be investigated. These variants could even be present at significant frequencies in populations missing or underrepresented in gnomAD.

Figure 5 illustrates the distribution of recalibrated mCSM-PPI2 ΔΔG predictions (ΔΔG^recal^) for all 475 possible ACE2 mutations at 25 ACE2 residues close to the Spike interface and the subset that are accessible via a single base change of the ACE2 coding sequence (these are more likely to be present in human populations than those requiring multiple substitutions). Most of these mutations are predicted to have only a slight effect on Spike binding, but there is a secondary mode below −1.0 kcal mol^−1^ with 126 mutations predicted to lead to strongly reduced binding. Fewer variants received high positive ΔΔG^recal^ scores: 17 had ΔΔG^recal^ > 1.0 kcal mol^−1^, a further 44 had ΔΔG^recal^ > 0.5 kcal mol^−1^ and an additional 70 had ΔΔG^recal^ > 0.2 kcal mol^−1^ (i.e., ΔΔG^recal^ > p.Lys26Arg). A similar pattern is observed for the 151 mutants corresponding to 172 single nucleotide variants of the ACE2 coding sequence (Figure 5B). These results suggest that random novel ACE2 missense variants at these loci can inhibit or enhance Spike binding, but are slightly more likely to be inhibitory, meaning that diversity at these positions might typically be beneficial and provide some resistance to infection. Notably, these recalibrated predictions do not display the same bias toward negative ΔΔG that the raw mCSM-PPI2 ΔΔG predictions did^23^ (Supplementary Figure 1), which may be another indicator of the improved quality of these recalibrated predictions.

**Figure 5.**
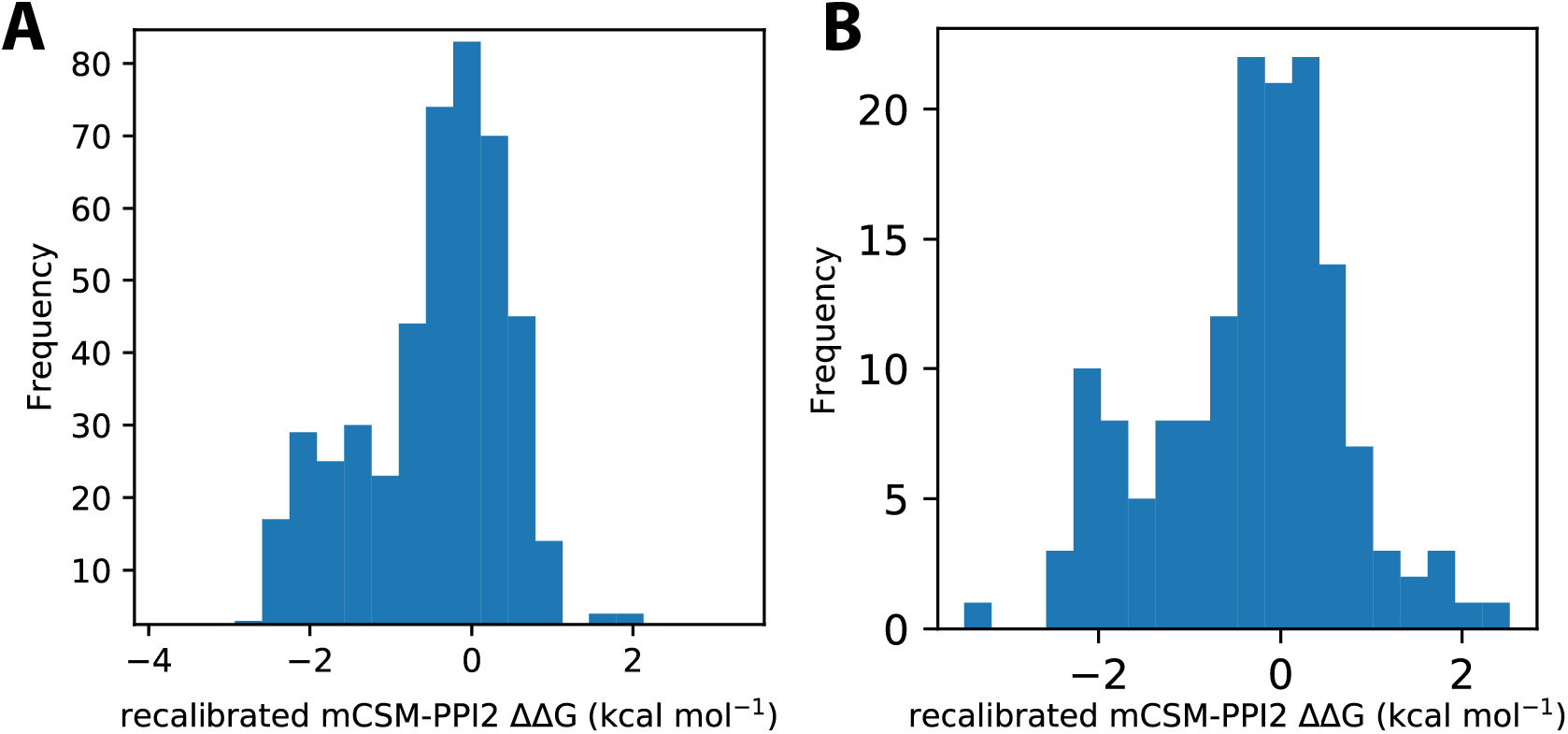
*A. Distribution of mCSM-PPI2^13^ predicted ΔΔG from in silico saturation mutagenesis of the ACE2-S interface in PDB 6vw1^11^. A. predicted ΔΔG for 475 mutations across 25 sites on ACE2 corresponding to the 23 residues within 5 Å of SARS-CoV-2 S plus Gly326 and Gly352. B. predicted ΔΔG for the subset of 151 mutations across these sites that are accessible via a single base change of the ACE2 coding sequence.*

Even though most of these variants are not reported in gnomAD, they may still occur within the populations represented, especially if they occur at frequencies that are poorly detected. With this in mind, we calculated allele frequencies for these mutations that are compatible with their absence from gnomAD (see Methods) in order to gain a better idea of how widespread the effects of these variants might be.

Table 3 presents estimated allele frequencies for novel variants that are predicted to inhibit and enhance Spike binding at varying ΔΔG^recal^ thresholds. Upper bounds for their joint frequencies assuming that they occur at lower frequencies than singleton variants suggest frequency bounds that range 1 in 1,000 to 1 in 2,000 variants per allele for inhibitory variants, and 1 in 1,500 to around 1 in 5,000 for enhancer variants. A second approach, which takes account of the empirical detection of rare variants in ACE2, yields frequencies that span from 1 in 6,250 to 1 in 12,195 for inhibitory variants, and 1 in 8,333 to 1 in 37,037 for enhancer variants. These estimates show that novel variants in ACE2 with any weak Spike affinity phenotype could plausibly be as common as 1 in 3,571 alleles (calculated as the sum of the highest inhibitor and enhancer prevalence’s), but the strongest affinity phenotypes are more likely to occur in frequency ranges akin to rare genetic diseases. It should be remembered that these values are calculated to be compatible with their absence from the gnomAD dataset, but it remains possible that some or all of these variants do not exist at all or that they are very common in one or more populations not represented in gnomAD.

**Table 3.** Estimated allele frequencies of potential novel variants in ACE2 with predicted Spike binding phenotypes. The estimate is calculated from the observed occurrence of rare variants in ACE2 in gnomAD24 (6.1 × 10-6) and the proportion of SNPs at the 25 residues that are predicted to modify Spike affinity at different thresholds (see Methods).

| $\Delta\Delta G^{\text{recal.}}$<br>(kcal mol<sup>-1</sup>) | N<br>missense | N<br>SNPs | Joint singleton<br>frequency<br>(NFE) | Joint estimated<br>frequency (NFE) | Heterozygotes / 100K | Hemizygotes / 100K |
| --- | --- | --- | --- | --- | --- | --- |
| <u>Predicted Spike inhibitory variants</u> |  |  |  |  |  |  |
| < -1.0 | 38 | 41 | 0.0005 | $8.2 \cdot 10^{-5}$ | 16.5 | 8.2 |
| < -0.5 | 55 | 59 | 0.0007 | 0.00012 | 23.8 | 11.9 |
| < -0.2 | 77 | 81 | 0.001 | 0.00016 | 33.0 | 16.5 |
| <u>Predicted Spike enhancer variants</u> |  |  |  |  |  |  |
| > 1.0 | 11 | 14 | 0.0002 | $2.7 \cdot 10^{-5}$ | 5.5 | 2.7 |
| > 0.5 | 24 | 30 | 0.0004 | $6.0 \cdot 10^{-5}$ | 11.9 | 6.0 |
| > 0.2 | 48 | 58 | 0.0007 | 0.00012 | 23.8 | 11.9 |

## 3 Discussion and conclusion

### 3.1 Impact of ACE2 affinity variants on SARS-CoV-2 infection

How far can affinity modulating variants influence SARS-CoV-2 infection? For inhibitory variants, host range specificity and mutagenesis studies^12^ highlight how some receptor mutations can provide complete protection from infection. We identified one variant (p.Asp355Asn) that abolished Spike binding altogether within our detection limits and a few others that reduced binding to varying extents. Since these variants were observed in population samples, some individuals carry ACE2 variants that could confer complete resistance to infection. Variants that reduce but do not eliminate binding may confer a degree of resistance proportional to the affinity reduction, or alternatively, there may be an affinity threshold that toggles cellular permissivity as in other enveloped viruses^33^. Detailed infectivity experiments with relevant ACE2 variants and multiple cell types are necessary to answer this question definitively.

It is less clear what effect affinity enhancing variants have on virulence, but there are indications that carriers may be at greater risk of infection and severe disease. Virus attachment proteins in enveloped viruses require a minimum receptor affinity that is proportional to the receptor surface density on the target cell to enable membrane fusion^33^. If this applies to SARS-CoV-2, ACE2 variants that enhance Spike binding could increase viral spreading in carriers due to increased cellular tropism. Greater viral spreading is associated with clinical deterioration^34^ and could cause increased infectiousness, akin to the enhanced transmissibility of SARS-CoV-2 vs. SARS-CoV, which is associated with increased viral loads in the upper respiratory tract^6,35^ and also correlates with enhanced receptor affinity^11^. The importance of tuned Spike-receptor affinities in SARS-CoV-2 may be enhanced because of the relatively low surface density of Spike on SARS-CoV-2 virions, which may lead to a weak avidity effect^36^. Anecdotal evidence is also provided by Spike RBD mutations, such as N501Y, that enhance ACE2 affinity^37^ and are associated with increased transmissibility^38^. Again, infectivity studies are needed but here the prevalence of p.Lys26Arg and p.Ser19Pro may allow the observation of an effect in future Covid-19 GWAS studies.

Some infectivity data for SARS-CoV-2 towards cells expressing ACE2 variants are available, including variants p.Ser19Pro and p.Lys26Arg^16^. Surprisingly, SARS-CoV-2 pseudotypes showed slightly decreased infectivity towards HEK293T cells expressing ACE2 p.Lys26Arg, and no difference compared to reference ACE2 was found for the susceptibility of cells expressing ACE2 p.Ser19Pro^16^. Although these results suggest that affinity is not directly proportional to infectivity in a single cell type, they do not exclude the possibility that these variants allow increased cellular tropism as discussed above. The same study also reported no effect on Spike affinity caused by these variants^16^, which is probably due to the low sensitivity of the cell based binding assay employed, since enhanced binding was also found in a different cell based assay^22^. Indeed, we are especially confident in the accuracy of our ΔΔG measurements since our results are consistent with published deep mutagenesis binding data^22^ (Supplementary Figure 2). In addition, another recent report agreed with our findings that the common ACE2 variants p.Ser19Pro and p.Lys26Arg enhanced affinity for Spike whilst p.Glu37Lys inhibits binding^17^.

Other considerations relevant to the effect of ACE2 Spike affinity variants on carriers include ACE2 carboxypeptidase activity^39^, which may be modulated by the specific binding affinity, the effect of hemizygosity and sex differences in Covid-19 outcomes, and the interplay of affinity variants and ACE2 expression levels. The importance of ACE2 expression was mentioned earlier but it is worth highlighting that it is known to partly determine the cellular specificity of SARS-CoV^40^ and was explored as a potential factor in COVID-19 susceptibility and severity, including the interaction between ACE2 variants and ACE2 stimulating drugs^41^. Also, heterozygotes express a proportion of ACE2 alleles whilst hemizygotes carry only a single ACE2 allele so that ACE2 Spike affinity variants ought to always show greater penetrance in hemizygotes, for better or worse. In contrast and since ACE2 escapes complete X-inactivation^20^, heterozygotes have the advantage that the more resistant ACE2 allele could become dominant in infected tissues due to selection over cellular infection cycles, gradually increasing the prevalence of the more Spike resistant ACE2 allele. Besides the possible benefit against infection, there could be other implications for heterozygotes depending on the persistence X-inactivation bias and the nature of any hitchhiking alleles.

### 3.2 Prevalence of ACE2-Spike affinity genotypes

Table 4 presents detailed allele frequencies from gnomAD^24^ for the ACE2 variants investigated in this work. The most prevalent ACE2 variant mediated Covid-19 phenotypes are likely to arise from the two relatively common variants found to enhance Spike binding, p.Ser19Pro and p.Lys26Arg. These variants were found in 3 in 1,000 individuals in the gnomAD African/African-American (AFR) samples and 7 in 1,000 North Western non-Finnish European samples (NW-NFE), respectively. p.Lys26Arg was also observed in other populations, including 1 in 1,000 AFR samples, adding further burden to this cohort. The high affinity displayed by ACE2 p.Ser19Pro is concerning in context with the reported disproportionate impact of Covid-19 on black and minority ethnic groups^30^. Further research to identify the impact of these variants on SARS-CoV-2 pathogenesis should be prioritised with urgency and existing and future GWAS should investigate these variants more closely with appropriate stratifications.

**Table 4:** *Detailed prevalence data from gnomAD^24^ for ACE2 variants reported in Table 1.*

| Variant | $\Delta\Delta G$ (kcal mol<sup>-1</sup>) | AFR | AMR | ASJ | EAS | FIN | NFE | SAS | OTH |
| --- | --- | --- | --- | --- | --- | --- | --- | --- | --- |
| p.Ser19Pro | 0.59 | 0.003 | - | - | - | - | - | - | 0.0002 |
| p.Lys26Arg | 0.31 | 0.001 | 0.003 | 0.01 | 0.00007 | 0.0005 | 0.006 | 0.001 | 0.003 |
| p.Thr27Ala | (-0.4) | - | 0.00007 | - | - | - | - | - | - |
| p.Glu35Lys | -0.29 | - | - | - | 0.0001 | - | 0.00001 | - | - |
| p.Glu37Lys | -1.25 | 0.0001 | - | - | - | 0.0003 | - | - | - |
| p.Phe40Leu | 0.14 | <sup>a</sup>0.0002 | <sup>b</sup>0.0001 | - | - | - | - | - | - |
| p.Met82Ile | (0.1) | 0.0003 | - | - | - | - | - | - | - |
| p.Gly326Glu | -0.67 | 0.0001 | - | - | - | - | - | - | - |
| p.Glu329Gly | -0.06 | - | - | - | - | - | 0.00007 | - | 0.0002 |
| p.Gly352Val | (-1.2) | - | - | - | - | - | 0.00001 | - | - |
| p.Asp355Asn | No binding | - | - | - | - | - | 0.00003 | - | - |
| p.Pro389His | 0.12 | - | 0.0002 | - | - | - | 0.00002 | - | - |
| Total frequency |  | 0.0047 | 0.00317 | 0.01 | 0.00017 | 0.0008 | 0.00614 | 0.001 | 0.0034 |
| Enhancers ($\geq 0.1$ kcal mol<sup>-1</sup>) | | 0.0045 | 0.0031 | 0.01 | 0.00007 | 0.0005 | 0.0062 | 0.001 | 0.0032 |
| Inhibitors ($\leq -0.1$ kcal mol<sup>-1</sup>) | | 0.0002 | 0.00007 | - | 0.0001 | 0.0003 | 0.00005 | - | - |

AFR: African/African-American, AMR: Latino/Admixed American, ASJ: Ashkenazi Jewish, EAS: East Asian, FIN: Finnish, NFE: non-Finnish European, SAS: South Asian, OTH: Other

The remaining gnomAD variants predicted to effect Spike binding are all very rare. The two other variants in our SPR series that also enhanced Spike binding have a joint frequency of around 3 in 10,000 in Latino/Admixed American (AMR) samples whilst amongst Spike inhibitory variants, p.Glu37Lys is the most prevalent occurring in 3 in 10,000 in Finnish samples and one additional African/African American sample. The other strongly inhibitory variants p.Gly352Val (predicted) and p.Asp355Asn are both doubletons observed only in non-Finnish Europeans, corresponding to an extremely low allele frequency of 3 in 100,000. Notably, the inhibitory variant p.Glu35Lys is the only variant predominant in East Asian samples

Our analysis of the effects of all possible missense mutations at the ACE2 Spike interface predicts many other potential novel variants that effect the Spike interaction, but even though a relatively high proportion of possible variants are predicted to effect Spike binding (up to 62%), their collective frequency could be as little as 2.8 in 10,000. These frequencies are comparable to those that might be expected for variants that cause rare diseases.

Although these data suggest that ACE2 alleles with large positive or negative ΔΔG are extremely rare it is important to note that allele frequencies show significant variation even between large variation datasets^42^. Moreover, it is known that local population allele frequencies can vary substantially from those reported in public datasets^43^ and it is therefore possible that some populations have risk or protective mutations at higher frequencies. It is also worth highlighting that the substantially increased Spike affinities of the two most common variants tested may indicate the presence of a past selective effect occurring twice independently, which may increase the possibility of other higher frequency variants amongst populations not well represented by gnomAD.

These frequencies might be useful for modellers to assess the role of rare ACE2 variants with Spike binding phenotypes on the epidemiology of Covid-19. For example, whether rare ACE2 variants that inhibit Spike binding contribute to the prevalence of asymptomatic carriers or if rare ACE2 variants with enhanced Spike binding can account for instances of particularly severe disease in the absence of typical comorbidities.

### 3.3 Conclusion

In summary, we report the binding affinities of 10 ACE2 variants with SARS-CoV-2 Spike RBD. We found that ACE2 p.Ser19Pro and p.Lys26Arg, two of the most common ACE2 missense variants in gnomAD^24^, substantially increase the affinity and are therefore possible risk factors for COVID-19. These variants have distinct distributions across the gnomAD cohorts and could therefore contribute population-specific risk. Two additional rare variants were identified that also enhanced Spike binding, although to a lesser extent. We confirmed our previous predictions that ACE2 p.Glu37Lys and p.Asp355Asn strongly inhibit Spike binding and are therefore potentially protective against COVID-19. We also identified two further rare variants that inhibited binding, including p.Gly326Glu, which we had previously incorrectly predicted would enhance binding. The SPR affinity data were used to recalibrate the mCSM-PPI2^13^ algorithm to provide improved affinity predictions for all possible ACE2 missense variants that interact with Spike and we estimated the prevalence of novel Spike affinity variants in ACE2. A key point in these burden assessments is the separate consideration of variants predicted to inhibit and enhance Spike affinity since these may have distinct phenotypes. Overall, p.Ser19Pro and p.Lys26Arg are still expected to have the most widespread effects, with the joint prevalence of the rare affinity variants substantially lower, but in all cases the possibility of higher prevalence in local or underrepresented populations remains and the penetrance of each variant will be a key factor. This work has applications in helping to prioritise experimental work into SARS-CoV-2 Spike human ACE2 recognition; developing genetic diagnostic risk profiling for COVID-19 susceptibility and severity, improving detection and interpretation in future COVID-19 genetic association studies and in understanding SARS-CoV-2 Spike evolution and host adaptation.

## Methods

### ACE2 and RBD constructs

The ACE2 construct was kindly provided by Ray Owens at the Oxford Protein Production Facility-UK. The RBD construct was kindly provided by Quentin Sattentau at the Sir William Dunn School of Pathology. ACE2 point mutations were added using Agilent QuikChange II XL Site-Directed Mutagenesis Kit following the manufactures instructions. The primers were designed using the Agilent QuikChange primer design web program.

### HEK293F suspension cell culture

Cells were grown in FreeStyle™ 293 Expression Medium (12338018) in a 37 °C incubator with 8% CO2 on a shaking platform at 130 rpm. Cells were passaged every 2-3 days with the suspension volume always kept below 33.3% of the total flask capacity. The cell density was kept between 0.5 and 2 million per ml.

### Transfection of HEK293F suspension cells

Cells were counted to check cell viability was above 95% and the density adjusted to 1.0 million per ml. For 100 ml transfection, 100 μl FreeStyle™ MAX Reagent (16447100) was mixed with 2 ml Opti-MEM (51985034) for 5 minutes. During this incubation 100 μg of expression plasmid was mixed with 2 ml Opti-MEM. For in situ biotinylation of ACE2 90 μg of expression plasmid was mixed with 10 μg of expression plasmid encoding the BirA enzyme. The DNA was then mixed with the MAX Reagent and incubated for 25 minutes before being added to the cell culture. For ACE2 biotinylation biotin was added to the cell culture at a final concentration of 50 μM. The culture was left for 5 days for protein expression to take place.

### Protein purification from HEK293F suspension cell supernatant

Cells were harvested by centrifugation and the supernatant collected and filtered through a 0.22 μm filter. Imidazole was added to a final concentration of 10 mM and PMSF added to a final concentration of 1 mM. 1 ml of Ni-NTA Agarose (30310) was added per 100 ml of supernatant and the mix was left on a rolling platform at 4 °C overnight. The supernatant mix was poured through a gravity flow column to collect the Ni-NTA Agarose. The Ni-NTA Agarose was washed 3 times with 25 ml of wash buffer (50 mM NaH2PO4, 300 mM NaCl and 20 mM imidazole at pH 8). The protein was eluted from the Ni-NTA Agarose with elution buffer (50 mM NaH2PO4, 300 mM NaCl and 250 mM imidazole at pH 8). The protein was concentrated, and buffer exchanged into size exclusion buffer (25 mM NaH2PO4, 150 mM NaCl at pH 7.5) using a protein concentrator with a 10,000 molecular weight cut-off. The protein was concentrated down to less than 500 μl before loading onto a Superdex 200 10/300 GL size exclusion column. Fractions corresponding to the desired peak were pooled and frozen at −80 °C. Samples from all observed peaks were analysed on an SDS-PAGE gel (Supplementary Figure 3).

### Surface plasmon resonance (SPR)

SARS-CoV-2 receptor binding domain binding to human extracellular ACE2 were analysed on a Biacore T200 instrument (GE Healthcare Life Sciences) at 37°C and a flow rate of 30 μl/min. Running buffer was HBS-EP (BR100669). Streptavidin was coupled to a CM5 sensor chip (29149603) using an amine coupling kit (BR100050) to near saturation, typically 10000-12000 response units (RU). Biotinylated ACE2 WT and variants were injected into the experimental flow cells (FC2–FC4) for different lengths of time to produce desired immobilisation levels (600–700 RU). FC1 was used as a reference and contained streptavidin only. Excess streptavidin was blocked with two 40 s injections of 250 μM biotin (Avidity).

Before RBD injections, the chip surface was conditioned with 8 injections of the running buffer. A dilution series of RBD was then injected simultaneously in all FCs. Buffer was injected after every 2 or 3 RBD injections. The lowest RBD concentration was injected at the beginning and at the end of each dilution series to ensure reproducibility. The length of all injections was 30 s, and dissociation was monitored from 180-300 s. Binding measured in FC1 was subtracted from the other three FCs. Additionally, all binding and dissociation data were double referenced using the closest buffer injections^44^. In all experiments, an ACE2-specific antibody (NOVUS Biologicals, AC384) was injected at 5 μg/ml for 10 minutes with the disassociation monitored for 10 minutes (Supplementary Figure 4). Only ACE2 T27R did not bind AC384 as expected but since this mutant displays RBD binding comparable to WT ACE2, this most likely indicates direct inhibition of AC384 binding rather than the presence of unfolded protein.

### SPR data fitting

Double referenced binding data was plotted and fit with GraphPad Prism (Supplementary Figure 5). To find the equilibrium K_D_ (dissociation constant) the association phase was fit with a One-phase association model and the plateau binding measurements were extracted and plotted against the corresponding concentration of RBD. This plot was then fit with a One-site specific binding model below and the value for the equilibrium K_D_ extracted.

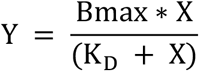

To convert K_D_ values to ΔG the equation below was used. Where R is the gas constant measured in cal mol^−1^K^−1^ and T is the temperature measured in K.

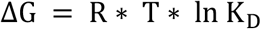

A ΔΔG value could then be found for each mutant by subtracting the ΔG of the WT from the ΔG of each mutant.

For ACE2 variant D355N binding was too poor to fit accurately therefore, an estimate for the lower limit for the K_D_ was calculated using the formula below. Where the “Maximum [RBD]” is the highest concentration of RBD flown over the surface, the “Binding at K_D_ (WT)” is the RU value at the equilibrium K_D_ for the WT protein on the same chip and the “Binding at maximum [RBD] (variant)” is the maximum RU value for the RBD binding D355N at the highest concentration. The estimated K_D_ lower limit could then be converted into ΔG and then ΔΔG using the same method above.

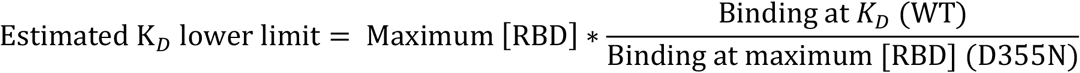

### Integration of structure, variant and mutagenesis data

The pyDRSASP suite^45^ was used to integrate 3D structure, population variant and mutagenesis assay data for analysis. Population variants from gnomAD v2^24^ were mapped to ACE2 with VarAlign^45^. Residue mappings were derived from the Ensembl VEP annotations present in the gnomAD VCF. In addition, we manually checked the gnomAD multi nucleotide polymorphisms (MNPs) data file and found no records for ACE2. The structure of chimeric SARS-CoV-2 Spike receptor binding domain in complex with human ACE2 (PDB ID: 6vw1)^11^ was downloaded from PDBe. Residue-residue contacts were calculated with ARPEGGIO^46^. These operations were run with our ProIntVar^45^ Python package, which processes all these data into conveniently accessible Pandas DataFrames. ACE2 Spike interface residues were defined as those with any interprotein interatomic contact (Supplementary Table 2).

### Prediction of missense variant effects on Spike – ACE2 interaction

The mCSM-PPI2^13^ web server was used to predict the effect of mutations on the SARS-CoV-2 Spike-ACE2 interface topology and binding affinity with the structure PDB ID: 6vw1^11^ according to our previous protocol^23^.

### Recalibrated mCSM-PPI2 predictions with SPR tested variants

The SPR determined ΔΔG (Kd-plateau) were regressed against the mCSM-PPI2 prediction, restricting the regression to the variants with negative predicted ΔΔG, with the *lm* function in R^47^. Recalibrated scores were calculated with the *predict.lm* function and applied to ACE2 variants within 10 angstroms of Spike.

### Enumerating possible ACE2 missense SNPs

The ACE2 gene (ENSG00000130234) was retrieved from Ensembl in Jalview^25^. Two identical CDS transcripts (ENST00000427411 and ENST00000252519) were found with the Get Cross-References command corresponding to ACE2 full-length proteins (ENSP00000389326 and ENSP00000252519). These correspond to the UniProt ACE2 sequence Q9BYF1. The CDS was saved in Fasta format and this was parsed in Python with Biopython. The CDS was broken into codons and all possible single base changes were enumerated and translated using the standard genetic code. This provided the set of amino acids accessible to each residue via a single base change.

### Estimated Prevalence of Novel Rare Variants in ACE2 with Spike Affinity Phenotypes

#### Upper bound from gnomAD singleton frequency (Minimum reportable frequency)

Upper bounds for the total prevalence of potential Spike affinity variants were calculated based on the conservative assumption that novel variants must occur at lower frequencies than the minimum reportable variant frequency (i.e., minimum singleton frequency) in gnomAD. The theoretical minimum reportable frequency is the allele frequency of a singleton variant at a site where all samples have been effectively called. For alleles on the X chromosome, the proportion of XX and XY samples is important since the number of alleles sequenced is 2*N*_female_ + *N*_male_. Practically, many reported singleton frequencies are greater than this theoretical minimum because at a given loci not all samples have sufficient sequence data quality to be called. Therefore, the minimum reportable frequency varies by genomic position, as well as population. Since gnomAD reports the allele number (AN) only for variant sites, we used the maximum observed allele number in ACE2. For example, the maximum allele number in ACE2 corresponding to non-Finnish European samples is 80,119 (AN_NFE = 80,119). This yields a minimum observed variant frequency in ACE2 amongst non-Finnish Europeans in gnomAD (v2 exomes) of 1/80,119 = 1.2 × 10^−5^, or 2.5 variants per 200,000 alleles. The total prevalence is then found by multiplying this value by the number of SNPs being considered.

#### Empirical detection rate of rare ACE2 variants and the affinity active ratio based estimate

Our second approach to estimate plausible frequencies of novel variants (*P*) was to project the empirical detection rate of rare ACE2 variants in gnomAD (*k*) onto the sites that correspond to the Spike interface (*n*), and then adjust this by the proportion of variants that are predicted to effect Spike binding (α) so that,

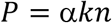

For example, there are 1,181 variant alleles in the 56,885 non-Finnish European cohort (AN_NFE = 80,119) arising from variants with AF < 0.01 in the 2,415 nucleotide ACE2 coding sequence (n.b. at this AF threshold, 90 % are missense). This corresponds to a variation rate *k* = 6.1 × 10^−6^ variants per nucleotide per allele. Projecting rate *k* onto the 25 ACE2 sites considered (*n* = 75 nucleotides) we find *kn* = 4.6 × 10^−4^ variants per allele, or 91.6 variants per 200,000 alleles. Note that this should be a conservative estimate of variability at these sites, since surface residues tend to be more variable than the core or other functional regions of the protein^48^, unless the site is under specific selection. The proportion of SNPs that are defined to have an affinity phenotype is dependent on the ΔΔG^recal^ threshold. For example, there are 38 substitutions corresponding to 41 single nucleotide variants out of the a possible 225 that are predicted to inhibit binding with ΔΔG^recal^ < −1.0 kcal mol^−1^ (Table 3) so that α = 41 / 225 = 0.18. Altogether this gives *P* = 8.2 10^−5^ for variants with ΔΔG^recal^ < −1.0 kcal mol^−1^. This calculation could be improved (e.g., to account for missense/ synonymous ratios in α) but in its current form is suitable to provide estimates that indicate plausible orders of magnitude of these variants’ prevalence as intended.

### Software

Jalview 2.11^25^ was used for interactive sequence data retrieval, sequence analysis, structure data analysis and figure generation. UCSF Chimera^26^ and PyMol^29^ were used for structure analysis and figure generation.

The pyDRSASP^45^ packages, comprising ProteoFAV, ProIntVar and VarAlign were used for data retrieval and analysis. Biopython was used to process sequence data.

Data analyses were coded in R and Python Jupyter Notebooks. Numpy, Pandas and Scipy were used for data analysis. Matplotlib, Seaborn and ggplot2 were used to plot data.

## Code availability

All code and analysis notebooks used in this study are available from the Barton Group public GitHub repository at https://github.com/bartongroup/covid19-ace2-variants.

## Data availability

This study employed several public datasets, which are available from their original sources and all necessary identifiers and accessions are provided in the methods. Data that was compiled and calculated as part of this study are available from the BioStudies database (https://www.ebi.ac.uk/biostudies) under accession S-BSST649.

## Acknowledgements

We thank Johannes Pettmann for help with protein expression and Anna Huhn for help with data analysis. We thank Dr Jim Procter for alerting us to the release of the structure of ACE2 in complex with SARS-CoV-2 S and his initial observations regarding variants at the ACE2-S interface. We thank the Dundee Research Computing team for supporting our IT infrastructure and remote working. This work was supported by Biotechnology and Biological Sciences Research Council Grants (BB/J019364/1 and BB/R014752/1) and Wellcome Trust Biomedical Resources Grant (101651/Z/13/Z). OD is supported by a Wellcome Trust Senior Fellowship in Basic Biomedical Sciences (207537/Z/17/Z828).

## Author contributions

SM performed the computational and population genetic analyses, compiled and analysed data, and wrote the first draft manuscript. MB conducted all wet lab experiments and compiled and analysed the associated data. GJB and SM conceived the research. All authors interpreted results and edited and reviewed the final manuscript.

## Author information

GJB and SM: Division of Computational Biology, School of Life Sciences, University of Dundee, Scotland, United Kingdom

MB, MK, OD and AvdW: Sir William Dunn School of Pathology, University of Oxford, Oxford, Oxfordshire, United Kingdom

## Extended Data

**Supplementary Table 1.**
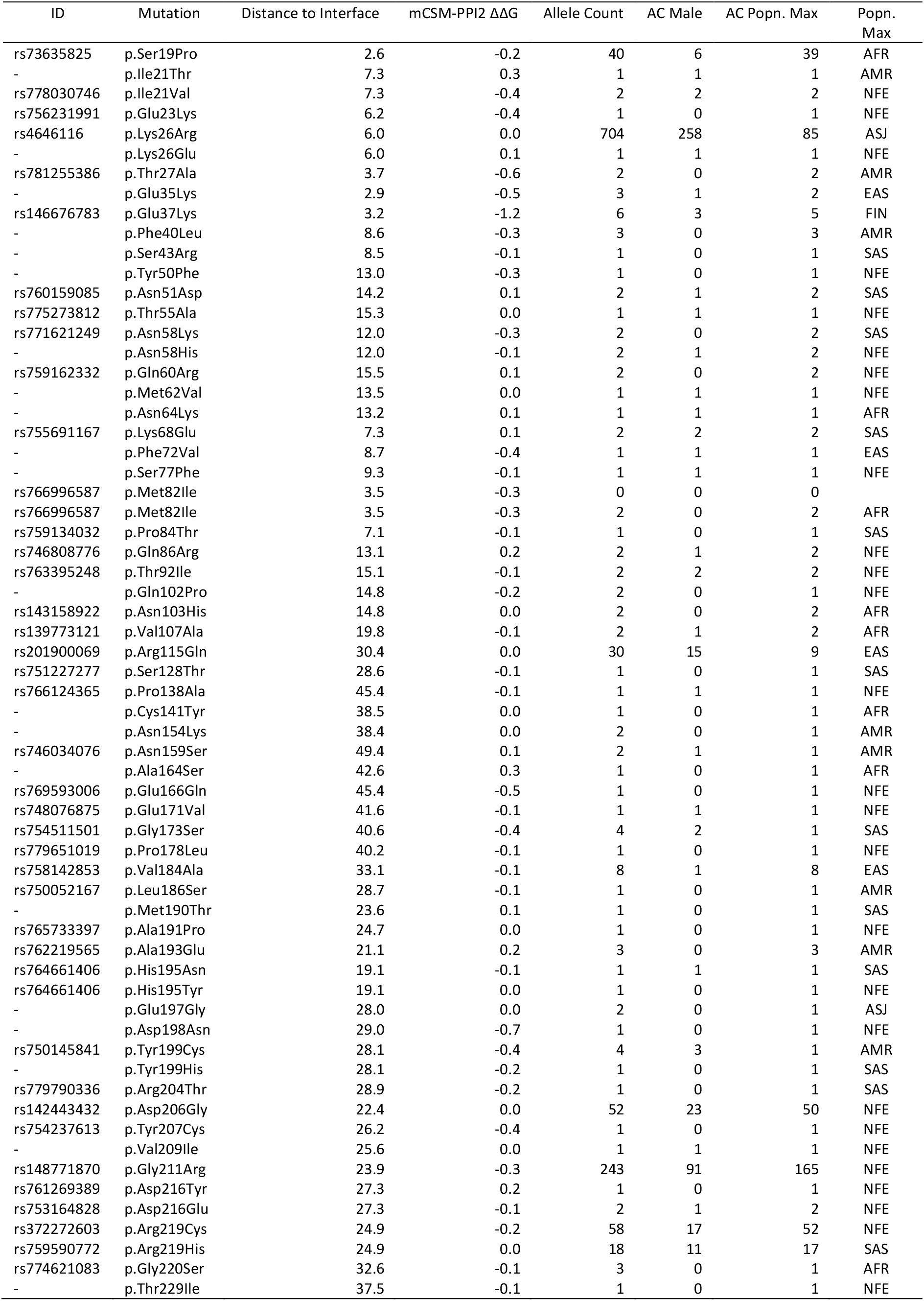
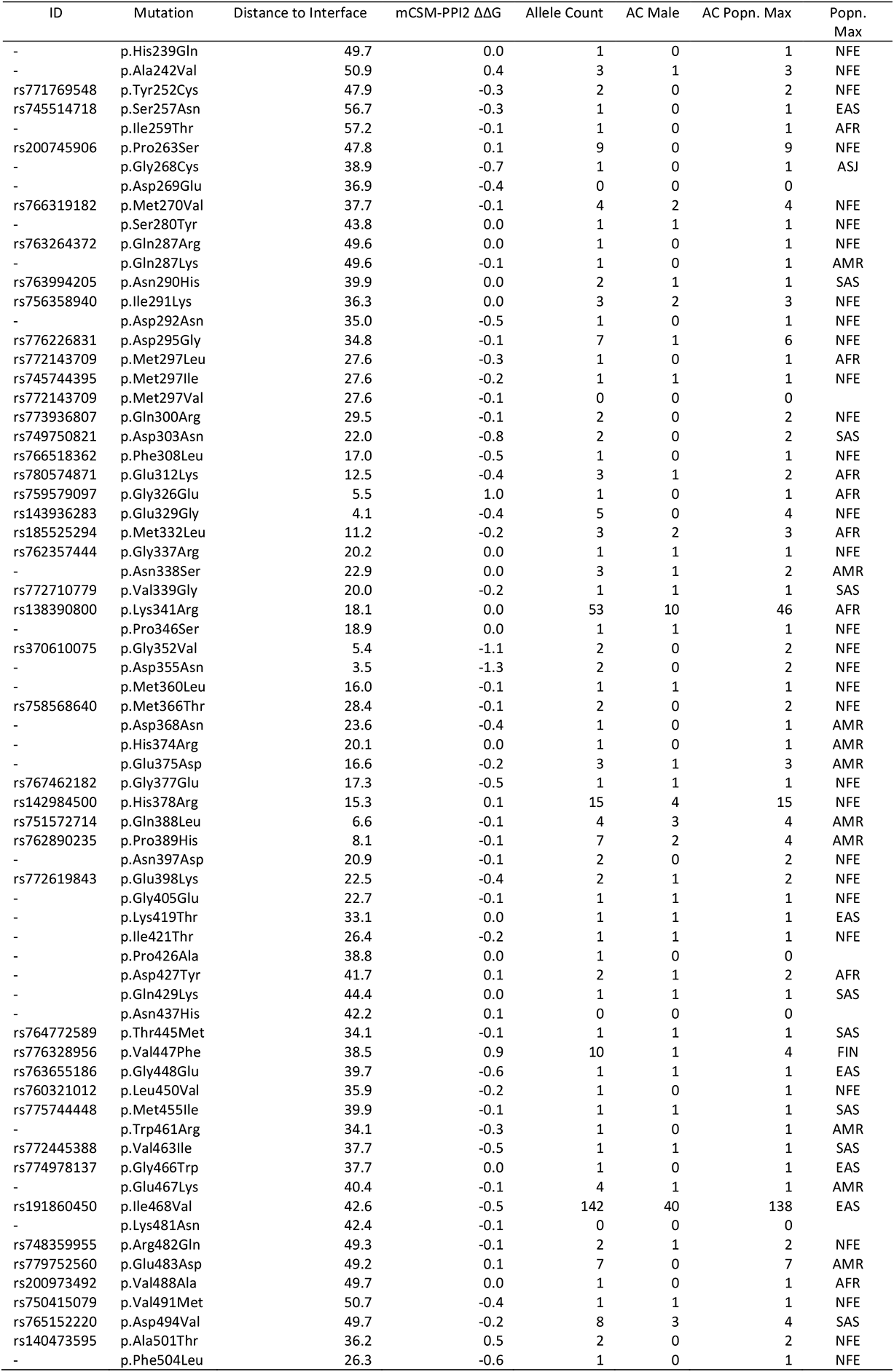
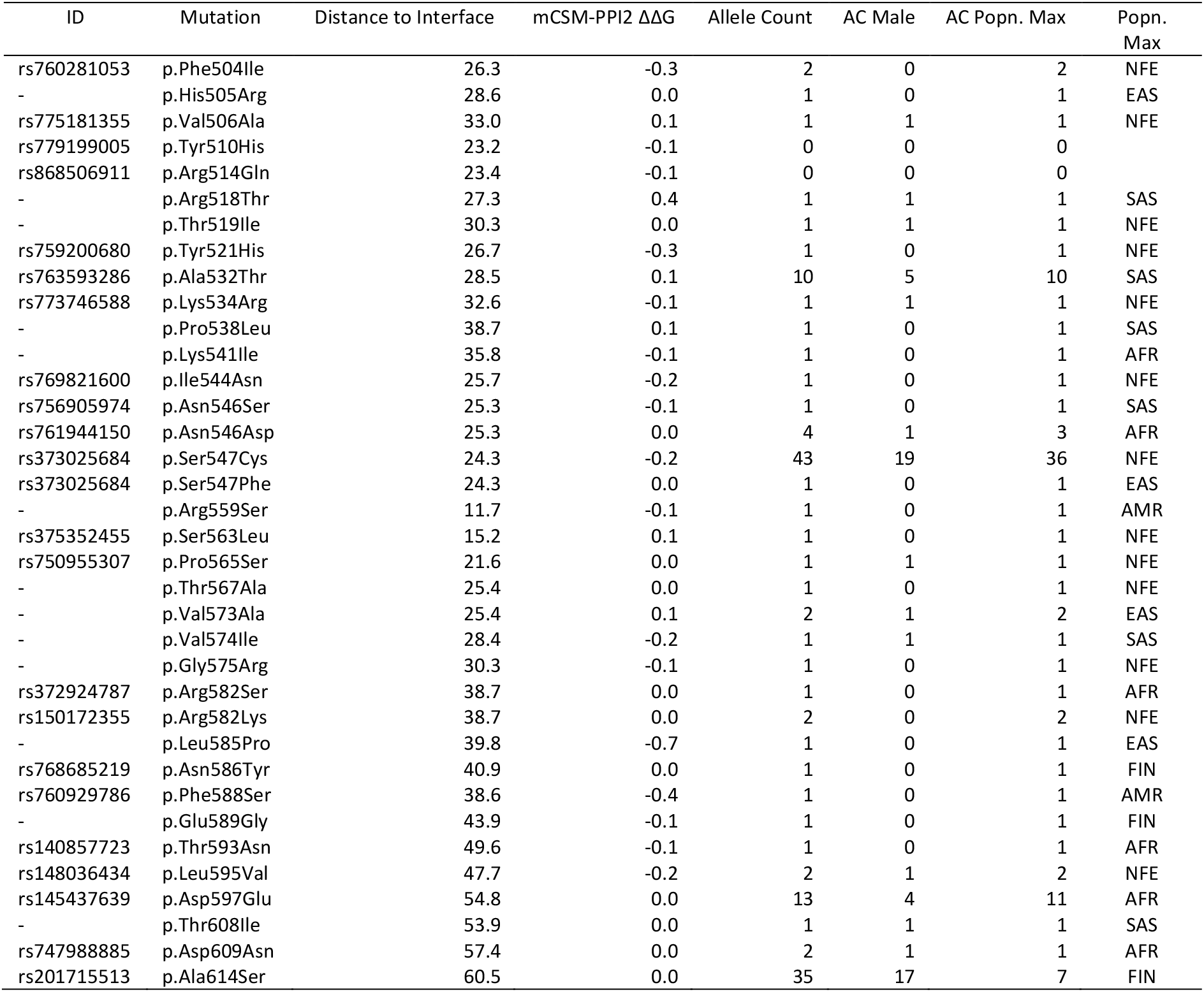
*mCSM-PPI2 predictions for all gnomAD ACE2 missense variants in residues resolved in PDB 6vw1.*

**Supplementary Table 2.**
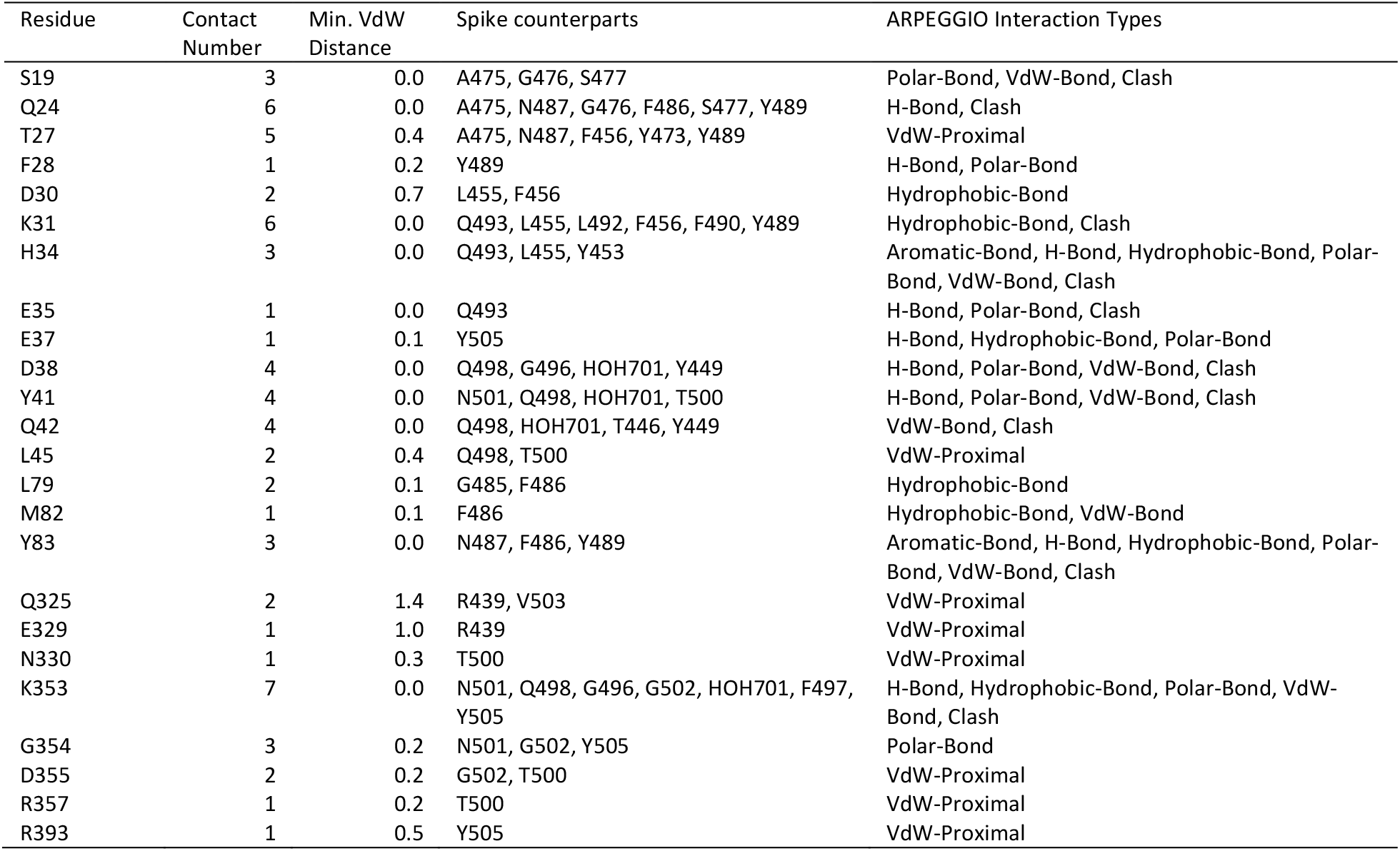
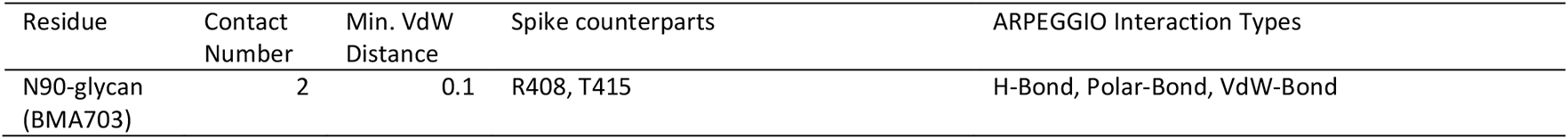
*Structural description of the ACE2 – Spike-protein interface.*

| Residue | Contact Number | Min. VdW Distance | Spike counterparts | ARPEGGIO Interaction Types |
| --- | --- | --- | --- | --- |
| S19 | 3 | 0.0 | A475, G476, S477 | Polar-Bond, VdW-Bond, Clash |
| Q24 | 6 | 0.0 | A475, N487, G476, F486, S477, Y489 | H-Bond, Clash |
| T27 | 5 | 0.4 | A475, N487, F456, Y473, Y489 | VdW-Proximal |
| F28 | 1 | 0.2 | Y489 | H-Bond, Polar-Bond |
| D30 | 2 | 0.7 | L455, F456 | Hydrophobic-Bond |
| K31 | 6 | 0.0 | Q493, L455, L492, F456, F490, Y489 | Hydrophobic-Bond, Clash |
| H34 | 3 | 0.0 | Q493, L455, Y453 | Aromatic-Bond, H-Bond, Hydrophobic-Bond, Polar-Bond, VdW-Bond, Clash |
| E35 | 1 | 0.0 | Q493 | H-Bond, Polar-Bond, Clash |
| E37 | 1 | 0.1 | Y505 | H-Bond, Hydrophobic-Bond, Polar-Bond |
| D38 | 4 | 0.0 | Q498, G496, HOH701, Y449 | H-Bond, Polar-Bond, VdW-Bond, Clash |
| Y41 | 4 | 0.0 | N501, Q498, HOH701, T500 | H-Bond, Polar-Bond, VdW-Bond, Clash |
| Q42 | 4 | 0.0 | Q498, HOH701, T446, Y449 | VdW-Bond, Clash |
| L45 | 2 | 0.4 | Q498, T500 | VdW-Proximal |
| L79 | 2 | 0.1 | G485, F486 | Hydrophobic-Bond |
| M82 | 1 | 0.1 | F486 | Hydrophobic-Bond, VdW-Bond |
| Y83 | 3 | 0.0 | N487, F486, Y489 | Aromatic-Bond, H-Bond, Hydrophobic-Bond, Polar-Bond, VdW-Bond, Clash |
| Q325 | 2 | 1.4 | R439, V503 | VdW-Proximal |
| E329 | 1 | 1.0 | R439 | VdW-Proximal |
| N330 | 1 | 0.3 | T500 | VdW-Proximal |
| K353 | 7 | 0.0 | N501, Q498, G496, G502, HOH701, F497, Y505 | H-Bond, Hydrophobic-Bond, Polar-Bond, VdW-Bond, Clash |
| G354 | 3 | 0.2 | N501, G502, Y505 | Polar-Bond |
| D355 | 2 | 0.2 | G502, T500 | VdW-Proximal |
| R357 | 1 | 0.2 | T500 | VdW-Proximal |
| R393 | 1 | 0.5 | Y505 | VdW-Proximal |
| N90-glycan (BMA703) | 2 | 0.1 | R408, T415 | H-Bond, Polar-Bond, VdW-Bond |

**Supplementary Figure 1.**
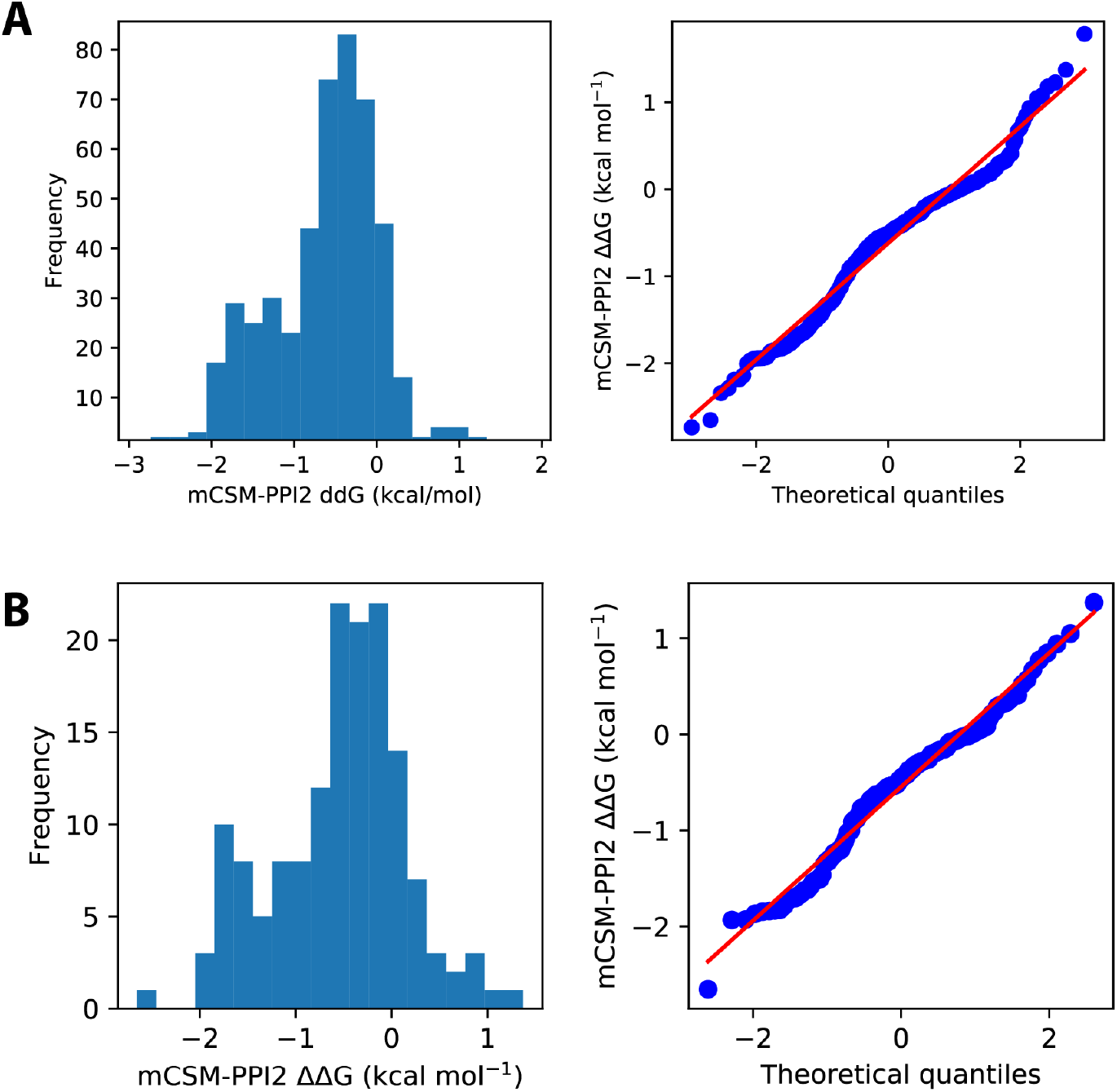
*A. Distribution of mCSM-PPI2^13^ predicted ΔΔG from in silico saturation mutagenesis of the ACE2-S interface in PDB 6vw1^11^. A. predicted ΔΔG for 475 mutations across 25 sites on ACE2 corresponding to the 23 residues within 5 Å of SARS-CoV-2 S plus Gly326 and Gly352. B. predicted ΔΔG for the subset of 151 mutations across these sites that are accessible via a single base change of the ACE2 coding sequence.*

**Supplementary Figure 2.**
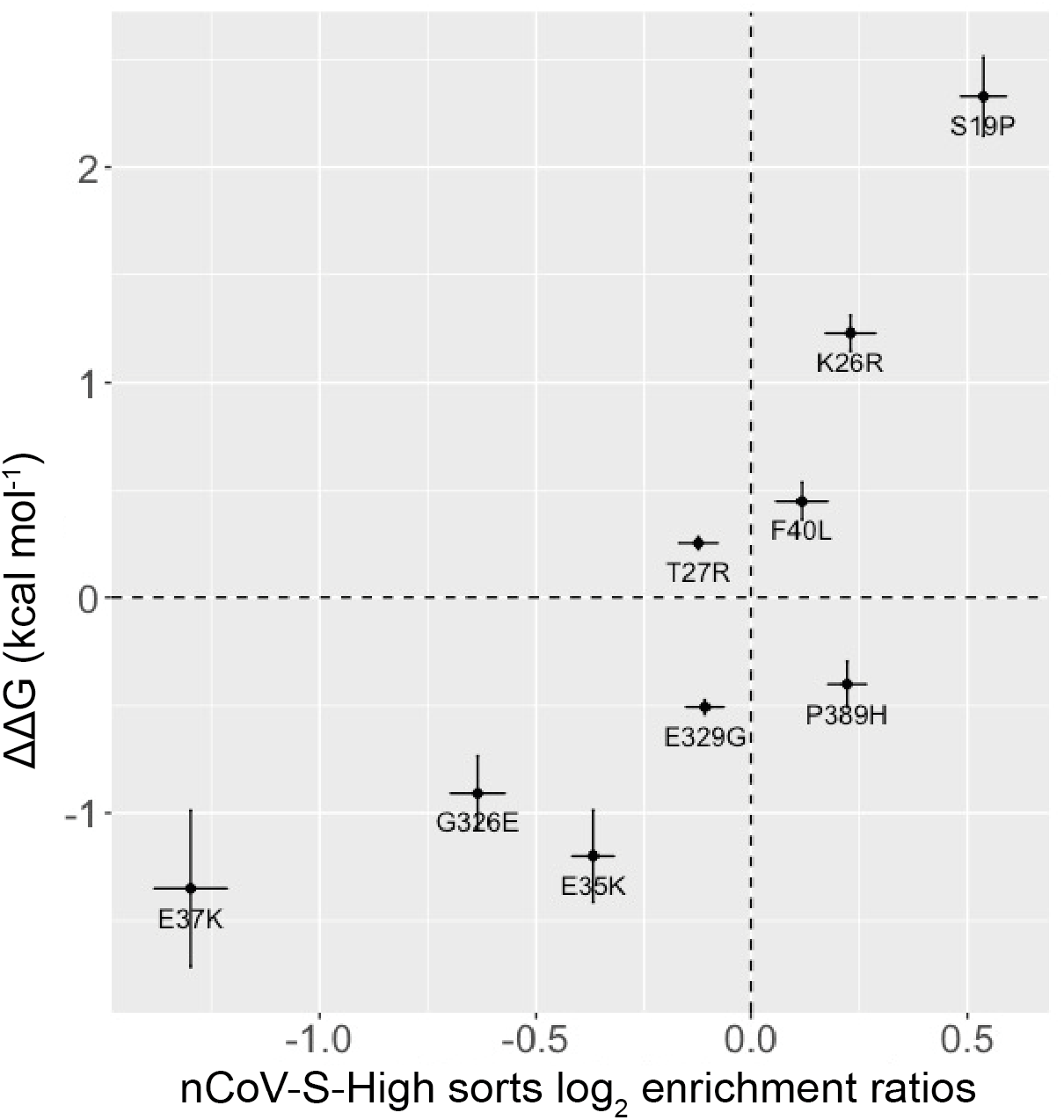
*Comparison of SPR ΔΔG with the log_2_ enrichment ratios from the nCoV-S-High sorts from Procko and co-workers^22^. Figure generated with R ggplot2.*

**Supplementary Figure 3.**
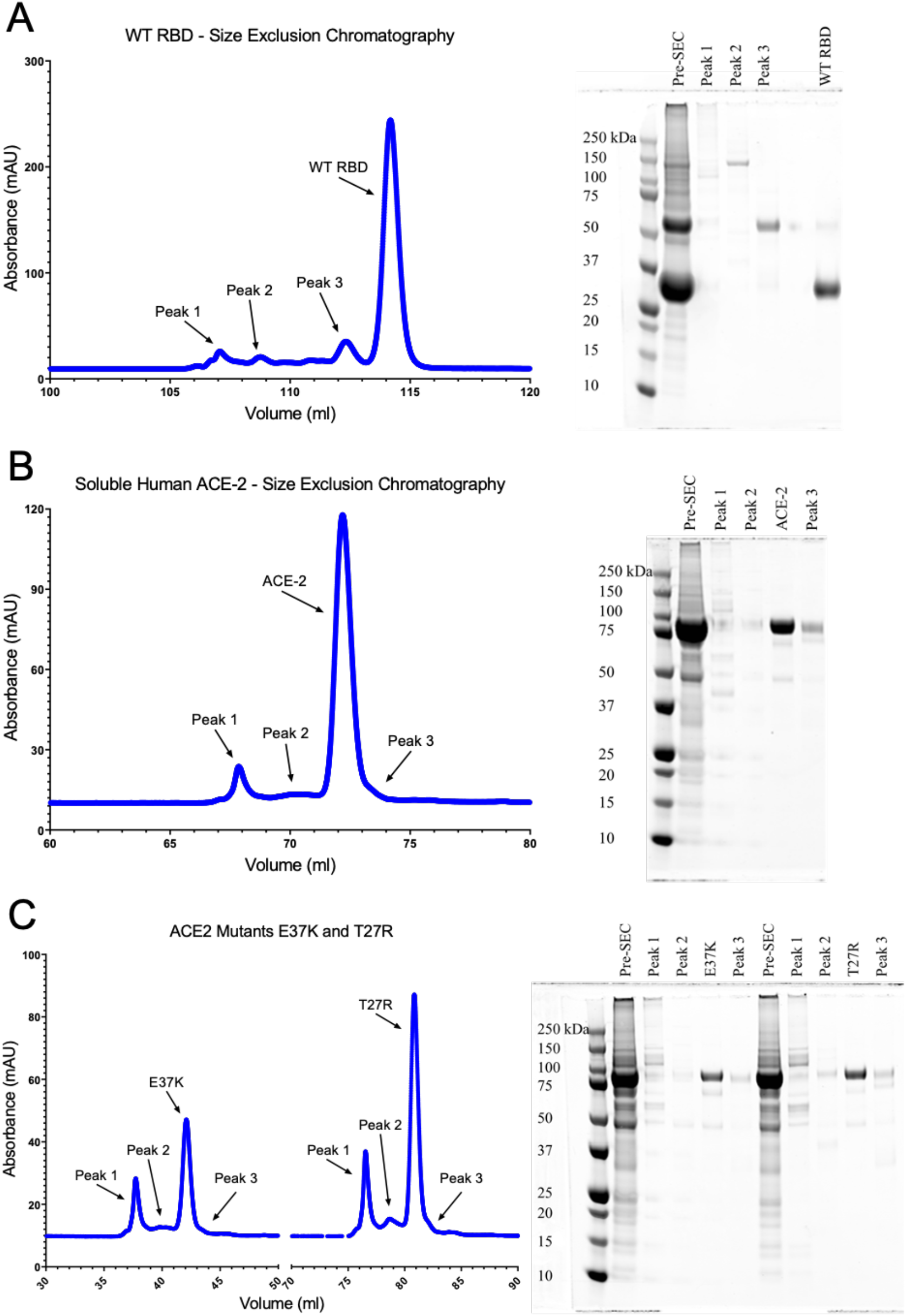
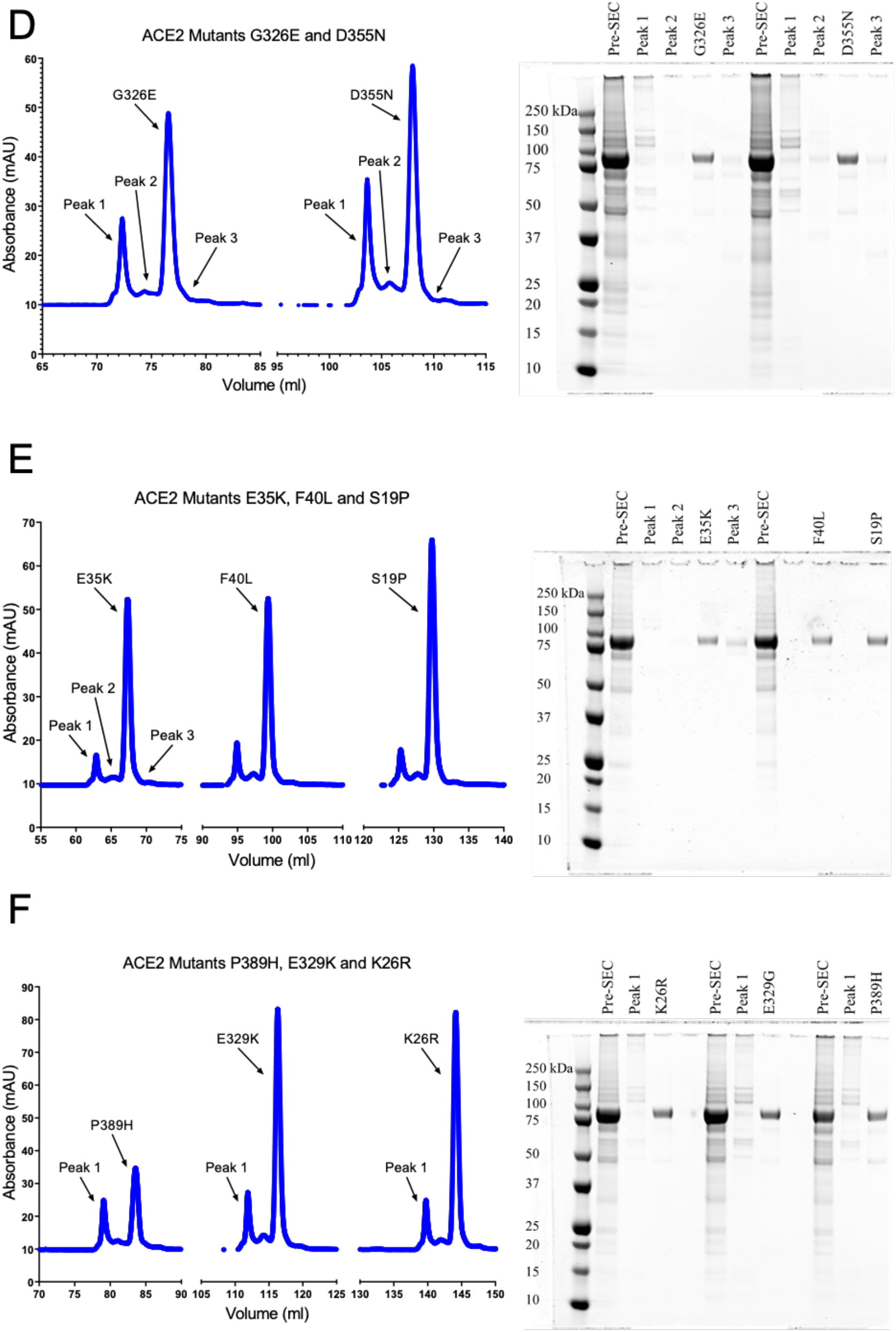
*Protein purification size exclusion chromatography and corresponding SDS-PAGE of labelled peaks. A. WT RBD, B. WT ACE2, (C) ACE2 mutants E37K and T27R, (continued next page) D. ACE2 mutants G326E and D355N, E. ACE2 mutants E35K, F40L and S19P, F. ACE2 mutants P329H, E329K and K26R. Each panel shows the l.*

**Supplementary Figure 4.**
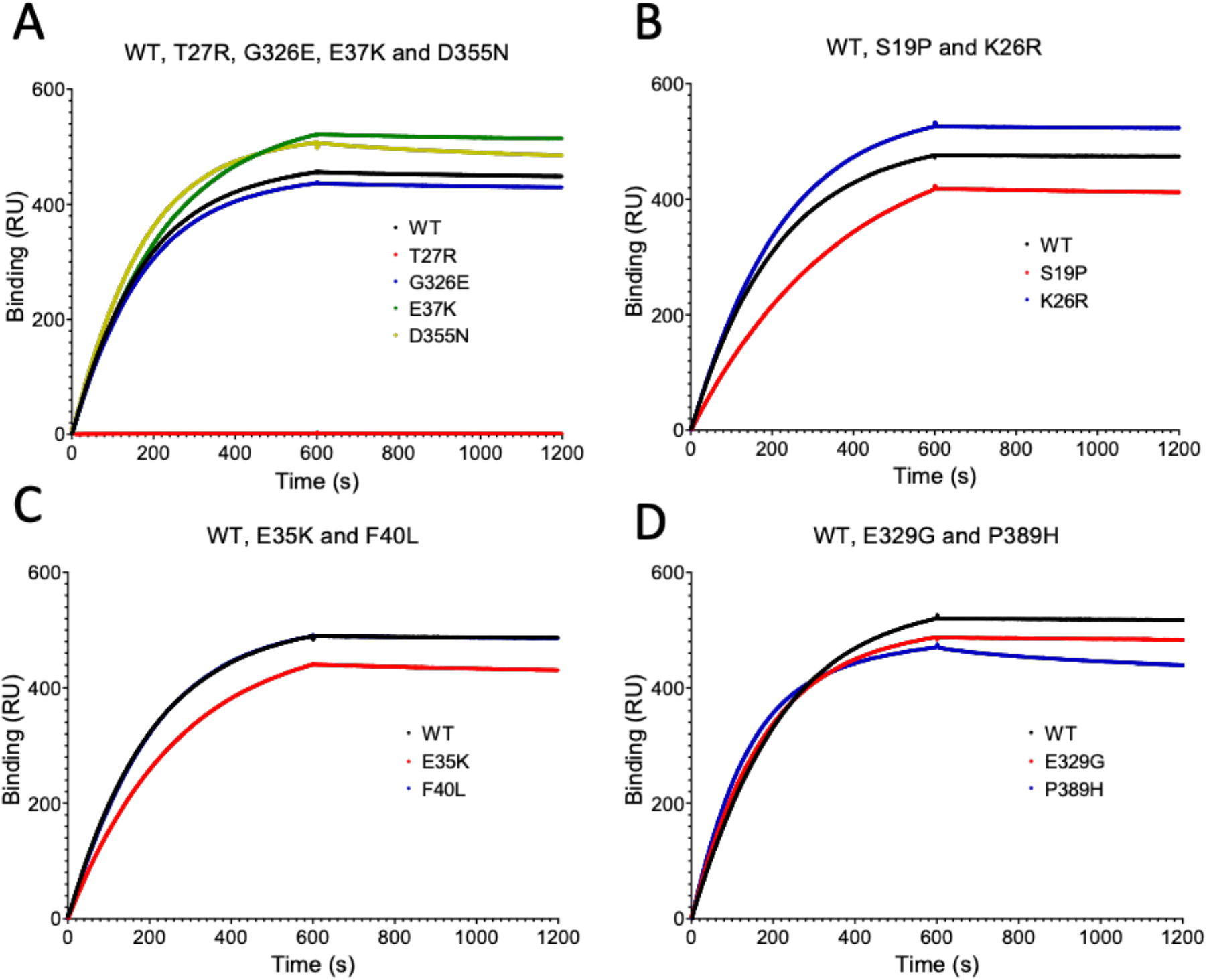
*Anti ACE2 Antibody (NOVUS AC384-NBP2-80038) binding to WT and 10 ACE2 variants. A. Anti ACE2 binding to WT and four ACE2 variants: T27R, G326E, E37K and D355N. B. Binding to WT, S19P and K26R. C. Binding to WT, E35K and F40L. D. Binding to WT, E329G and P389H. Anti ACE2 was injected for 600 s association and buffer for a further 600 s for dissociation.*

**Supplementary Figure 5.**
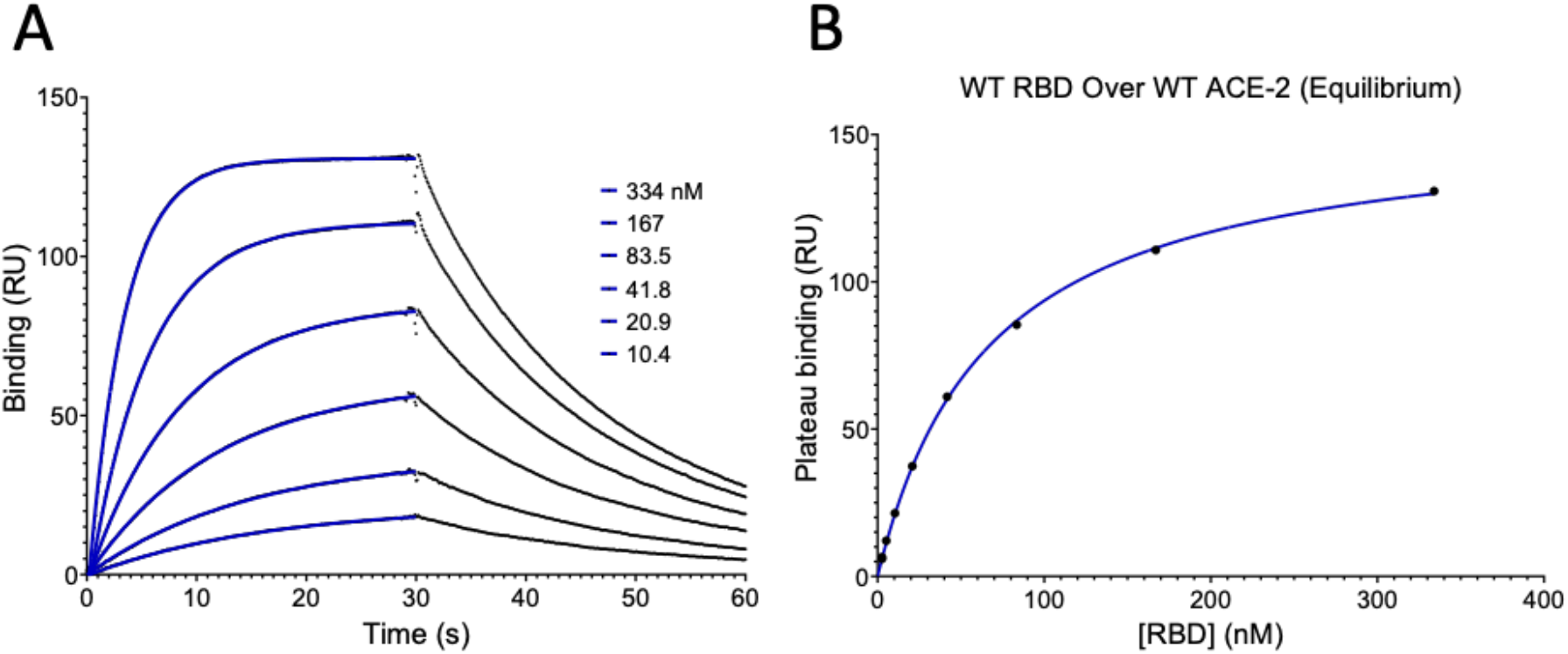
*WT RBD binding WT ACE2 plus equilibrium K_D_. A. Representative binding trace for WT RBD binding immobilised WT ACE2. Injections start from lowest to highest concentration of RBD and fits are shown in blue. B. Plateau binding values from A are plotted against the concentration of RBD injected and the equilibrium K_D_ is extracted from the fit shown in blue.*

